# Bystander memory-phenotype conventional CD4^+^ T cells exacerbating autoimmune neuroinflammation

**DOI:** 10.1101/2022.06.17.496529

**Authors:** Min-Zi Cho, Hong-Gyun Lee, Jae-Won Yoon, Gil-Ran Kim, Ja-Hyun Koo, Reshma Taneja, Brian T. Edelson, You Jeong Lee, Je-Min Choi

## Abstract

Memory-phenotype (MP) CD4^+^ T cells are a substantial population of conventional T cells that exist in steady-state mice, and their immunologic functions in autoimmune disease have not yet been studied. In this work, we unveil a unique phenotype of MP CD4^+^ T cells by analyzing single-cell transcriptomics and T cell receptor (TCR) repertoires. We found that steady-state MP CD4^+^ T cells exist regardless of germ and food-antigen which are composed of heterogenous effector subpopulations. Distinct subpopulations of MP CD4^+^ T cells are specifically activated by IL-1 family cytokines and STAT activators, revealing that the cells have TCR-independent effector functions. Especially, CCR6^high^ MP CD4^+^ T cells are major responders to IL-1β and IL-23 without MOG_35-55_ antigen reactivity, which gives them pathogenic-Th17 characteristics and allows them to contribute to autoimmune encephalomyelitis. We identified Bhlhe40 in CCR6^high^ MP CD4^+^ T cells drives the expression of GM-CSF, contributing to CNS pathology in experimental autoimmune encephalomyelitis. Collectively, our findings reveal heterogeneity of MP CD4^+^ T cells that can contribute to autoimmune neuroinflammation in bystander manner synergistically with antigen-specific T cells.

## Introduction

The immunological memory of antigen-specific T cells enables faster and more potent responses upon re-exposure to a previously encountered antigen, providing long-lasting immunity (Kaech & Cui, 2012). Although a substantial population of memory-phenotype (MP) conventional T cells exists in steady state mice unexposed to foreign antigens, it has been reported that MP T cells are formed before birth in humans and exist in germ-free (GF) and antigen-free (AF) conditioned mice (Byrne *et al*, 1994; Dobber *et al*, 1992; Haluszczak *et al*, 2009; Szabolcs *et al*, 2003).

After undergoing homeostatic proliferation in a lymphopenic environment, naïve T cells acquire phenotypical, functional, and gene expression–like antigen-specific memory and become MP T cells (Cho *et al*, 2000; Ernst *et al*, 1999; Goldrath & Bevan, 1999; Goldrath *et al*, 2004). T cell receptor (TCR) and CD28 signaling seem to be required for the conversion of naïve CD4^+^ T cells into MP T cells. MP CD4^+^ T cells express CD5, a marker with high affinity to self-antigen, and resist infection by mediating a Th1-like immune response without antigen stimulation (Kawabe *et al*, 2017). A previous study confirmed that antigen-non-specific MP CD4^+^ T cells proliferated more than Lymphocytic choriomeningitis virus (LCMV)-specific T cells in an LCMV infection model. Indeed, treatment with an anti-MHCII antibody did not inhibit the proliferation of MP CD4^+^ T cells, suggesting that they play a bystander role against infection (Younes *et al*, 2011).

Unlike MP CD4^+^ T cells, MP CD8^+^ T cells have been well studied for their antigen-specific and bystander functions. MP CD8^+^ T cells can rapidly produce IFN-γ upon stimulation with IL-12 and IL-18 and without cognate antigen stimulation (Sosinowski *et al*, 2013; White *et al*, 2017). In particular, MP CD8^+^ T cells are called virtual memory. They can have specific reactions to certain antigens without previous exposure (Akue *et al*, 2012; Haluszczak *et al*., 2009; Hamilton *et al*, 2006; Sosinowski *et al*., 2013; White *et al*, 2016) and increase NKG2D and granzyme B expression upon IL-12, IL-18, and IL-15 stimulation, producing a bystander killing role against infection (Chu *et al*, 2013; Lertmemongkolchai *et al*, 2001; White *et al*., 2016). In addition, MP CD8^+^ T cells can play an antigen-specific protection role in *Listeria monocytogenes*, Herpes simplex virus (HSV), and *vaccinia* infections (Haluszczak *et al*., 2009; Hamilton *et al*., 2006; Lee *et al*, 2013a; Sosinowski *et al*., 2013). Also, MP CD8^+^ T cells have high affinity to self-antigens, so they can break peripheral tolerance with MP CD4^+^ T cells and develop autoimmune diabetes (King *et al*, 2004; Le Saout *et al*, 2008). Overall, previous studies have shown the function of MP CD8^+^ T cells in disease, but the role of MP CD4^+^ T cells in autoimmune disease has not yet been studied.

Most previous studies have focused on the role of autoantigen-specific T cells in both humans and mice to understand autoimmune diseases and find therapeutic drugs to regulate antigen-specific T cells (Serra & Santamaria, 2019). Interestingly, antigen-non-specific T cells, including myelin oligodendrocyte glycoprotein (MOG) tetramer–negative CD4^+^ Th17 cells, also infiltrate the Central nervous system (CNS) in significant proportions and exacerbate experimental autoimmune encephalomyelitis (EAE) pathogenesis (Jones *et al*, 2003; Lee *et al*, 2019; Lees *et al*, 2010; Lin *et al*, 2016). In support, myelin-specific T cells were found to be similar in Multiple sclerosis (MS) patients and healthy controls (Hellings *et al*, 2001; Hemmer *et al*, 1997). Bystander-activated T cells and Epstein-Barr virus–specific CD8^+^ T cells are clonally expanded and correlate with disease pathogenesis in the joints of chronic inflammatory arthritis and Sjogren’s syndrome patients (Brennan *et al*, 2008; Kobayashi *et al*, 2004; Tan *et al*, 2000), raising questions about the role of antigen-nonrelated T cells in autoimmune disease.

In this study, we hypothesized that MP conventional CD4^+^ T cells could be encephalitogenic bystander cells during the development of autoimmune neuroinflammatory disease. First, we examined the heterogeneity of MP CD4^+^ T cells using single cell RNA-sequencing (scRNA-seq) and TCR sequencing analyses. We found distinct subpopulations of Th1-, Th17-, Treg-, and Tfh-like cells among MP CD4^+^ T cells. Those cells respond to IL-1 family cytokines and STAT activators, even without TCR stimulation. We further found that CCR6 ^high^ MP CD4^+^ T cells are the major responders to IL-1β and IL-23, expressing pathogenic signature genes in a bystander manner and thereby contributing to the development of MOG antigen–specific T cell–derived EAE. In this context, Bhlhe40 regulates the production of GM-CSF in CCR6^high^ MP CD4^+^ T cells, exacerbating EAE pathogenesis. Our findings indicate the pathogenic role of antigen-unrelated MP conventional CD4^+^ T cells along with antigen-specific T cells during autoimmune neuroinflammatory disease.

## Results

### Single cell RNA-sequencing identifies distinct effector-like phenotypes in steady-state MP conventional CD4^+^ T cells

Steady-state, unprimed specific-pathogen-free (SPF)-housed mice have a significant proportion of MP CD4^+^ T cells (TCRβ^+^CD1d tetramer^-^CD8^-^CD4^+^CD25-CD44^high^CD62L^low^) in their spleen, thymus, inguinal lymph nodes (iLNs), mesenteric lymph nodes (mLNs), Peyer’s patches (PPs), and lung tissues (Fig. S1A and Fig. 1A). The proportion and number of CD4^+^ T cells increased with age in all tissues (Fig. S1). To identify and focus on the characteristics of MP CD4^+^ T cells, we used fluorescence-activated cell sorting (FACS) to sort MP CD4^+^ T cells from the spleens of 10-week-old C57BL/6 mice and performed scRNA-seq with paired V(D)J sequencing of the T cell receptor (Fig. S2A to D). Also, we generated a pipeline to filter out PLZF and TCR V_α_14-J_α_18 (TRAV11-TRAJ18) expressing cells, a well-known key transcription factor for the development of NKT and MAIT cells (Koay *et al*, 2016; Kovalovsky *et al*, 2008; Savage *et al*, 2008). An unbiased clustering analysis revealed clearly distinct effector T cell–like subpopulations (Fig. 1B) and differentially expressed genes (DEGs) defining each cluster (Fig. 1C and D). A Gene Ontology (GO)/Kyoto Encyclopedia of Genes and Genomes (KEGG) analysis of each cluster indicated that clusters 3 and 5 were Th1- and Th17-like populations, respectively, with significance in “chemokine-mediated signaling pathway” and “cellular response to interleukin-1” (Fig. S3). In addition, lineage-specific transcription factors and chemokine receptors were localized in each cluster (Fig. 1E and Fig. S4), suggesting that MP CD4^+^ T cells are composed of Th1-, Th17-, Tfh-, and Treg-like populations. To examine whether the subpopulations of steady-state MP CD4^+^ T cells depended on germ or food antigens, we analyzed the proportion of CD44^high^CD62L^low^ MP CD4^+^ T cells in the tissues of SPF-, GF-, and AF-housed mice using flow cytometry. The proportions of total MP CD4^+^ T cells were almost identical in the spleen and other tissues except the mLN and PP (Fig. 1F, Fig. S5A and S5B), indicating that the generation of splenic MP CD4^+^ T cells is not affected by the microbiome or food antigen stimulation. Our scRNA-seq analysis further confirmed that the SPF- and GF-housed mice had identical proportions of distinct subpopulations of splenic MP CD4^+^ T cells (Fig. 1G and 1H), but the proportion of Th17-like MP cells in the mLNs of GF-housed mice was much lower than that in the mLNs of SPF-housed mice (Fig. 1I and 1J). In support, the proportion of CCR6^high^ or RORγt^+^ MP CD4^+^ T cells in the mLNs of GF-housed mice was lower than that in SPF-housed mice, suggesting that the generation of gut MP CD4^+^ T cells is specifically dependent on germs (Fig. S6). The TCR clonal diversity of MP CD4^+^ T cells in GF-housed mice, especially the Th17-like population in the mLNs, was reduced compared with SPF-housed mice, though comparable diversity was observed in the spleen (Fig. 1K and 1L). Together, these data indicate that steady-state splenic MP CD4^+^ T cells contain heterogeneous subpopulations of Th1-, Th17-, Tfh-, and Treg-like cells that express effector molecules and exist independently of the gut microbiome and food antigens.

**Figure 1.**
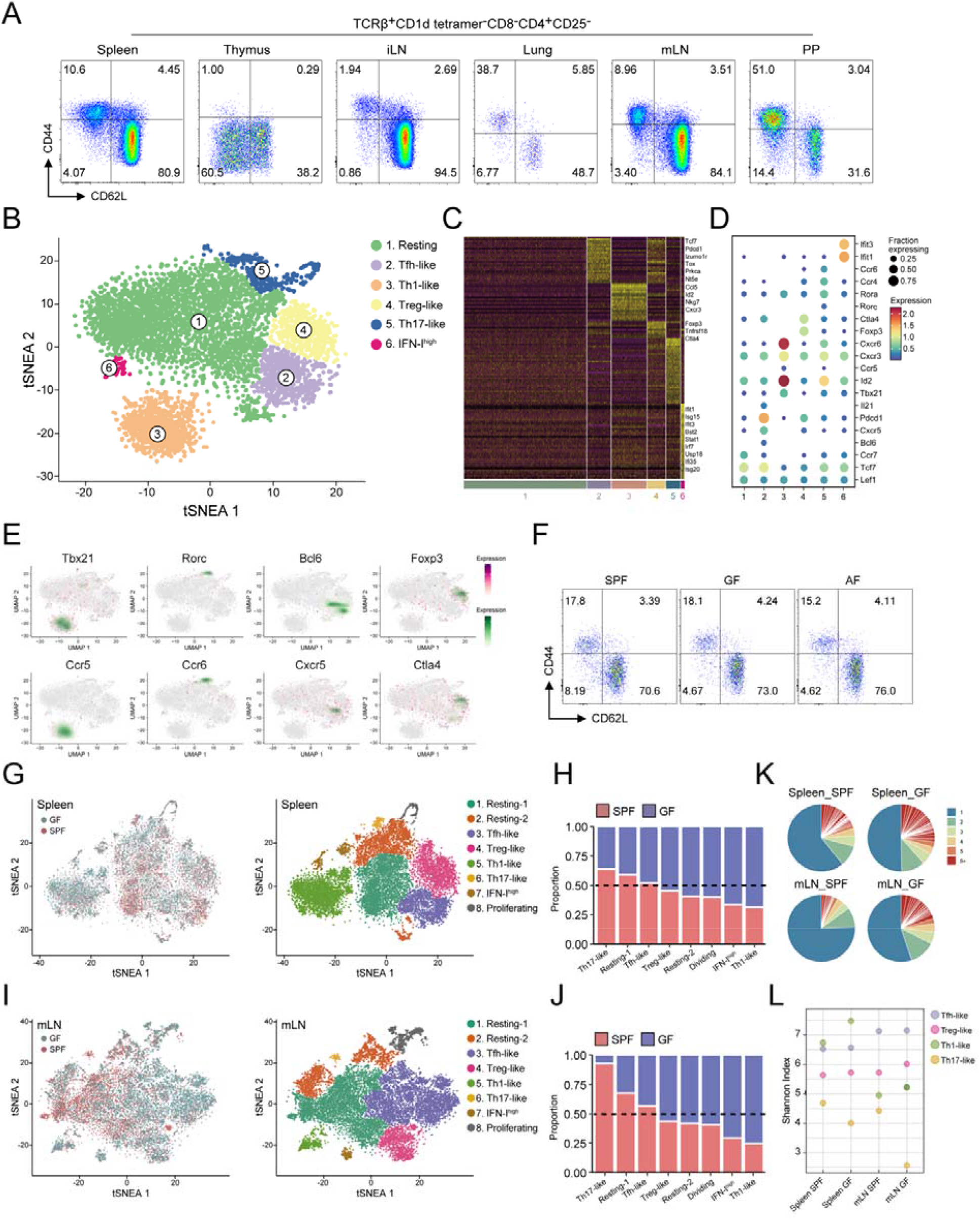
Single cell RNA-sequencing identifies distinct effector-like phenotypes in steady-state MP CD4^+^ T cells. (A) Representative flow cytometry plots showing the population of memory-phenotype (MP) CD4^+^ T cells (TCRβ^+^CD1d tetramer^-^CD8^-^CD4^+^CD25^-^CD62L^low^CD44^high^) in murine spleens, thymuses, inguinal lymph nodes (iLNs), lungs, mesenteric lymph nodes (mLNs), and Peyer’s patches (PP). (B) tSNE plot of splenic MP CD4^+^ T cells isolated from 10-week-old specific-pathogen-free (SPF) mice. (C) Heatmap of differentially expressed transcripts in splenic MP CD4^+^ T cells. (D) Expression of selected genes used to define MP CD4^+^ T cell clusters. (E) Differential expression of transcription factors and chemokine receptors from splenic MP CD4^+^ T cell clusters. (F) Flow cytometry plots of splenic MP CD4^+^ T cells (TCRβ^+^CD1d tetramer^-^CD8^-^CD4^+^CD25^-^CD62L^low^CD44^high^) from SPF-, GF-, and AF-mice. (G) tSNE plot and (h) proportion of integrated splenic MP CD4^+^ T cells isolated from 10-week-old SPF- and GF-mice. (I) tSNE plot and (J) proportion of integrated mLN MP CD4^+^ T cells isolated from SPF- and GF-mice. (K) Pie chart representing the clonal size distribution of MP CD4^+^ T cells. (L) Diversity of the TCR repertoire in MP CD4^+^ T cell subsets from SPF- and GF-mice.

### Phenotypic differences of MP CD4^+^ T cells after exposure to IL-1 family and STAT activating cytokines

To determine the effector functions of MP CD4^+^ T cells to respond to various cytokines, we performed scRNA-seq of MP CD4^+^ T cells cultured in conditioned medium with IL-12/IL-18, IL-25/IL-33, and IL-1β/IL-23 cytokines (Fig. 2A and Fig. S2E). Additionally, IL-7 was added to the culture medium for T-cell survival and maintenance (Kondrack *et al*, 2003). The merged Uniform Manifold Approximation and Projection (UMAP) plot shows phenotypic characteristics of Th1-, Th2-, Th17-, and Treg-like subpopulations (Fig. 2B and C), which express lineage specific genes (Fig. 2D). Each cytokine set induced distinct subpopulations localized exclusively in the plot (Fig. 2E). We further confirmed the expression of selected gene sets related to the Th1, Th2, and Th17 lineages in each bystander-activated condition, which shows that type 1, 2, and 3 cytokines can upregulate lineage-specific genes in MP CD4^+^ T cells (Fig. 2F and G). A single-cell regulatory network inference and clustering (SCENIC) analysis in each cytokine condition revealed a significant level of common or specific transcription factor activity (Fig. 2H). In addition, an Ingenuity pathway analysis (IPA) showed the predicted transcriptional regulators in each condition and speculated that the target transcription factors included Bhlhe40, which can be a potent regulator of bystander activation in MP CD4^+^ T cells (Fig. 2I). To evaluate the effector functions of MP CD4^+^ T cells, we treated MP CD4^+^ T cells with those cytokines and/or anti-CD3/anti-CD28 (Fig. 2J and 2K). Consistently, the MP CD4^+^ T cells responded to IL-12/IL-18, IL-25/IL-33, and IL-1β/IL-23 cytokines without TCR stimulation, and T-bet^+^ IFN-γ^+^, GATA3^+^IL-13^+^, and RORγt^+^IL-17^+^ cells increased compared to TCR-stimulated condition (Fig. 2J). However, TNF-α was produced only in the presence of TCR stimulation and IFN-γ in the culture supernatant from IL-12/IL-18-conditioned MP CD4^+^ T cells, and IL-4 was also significantly detected with TCR stimulation but not in the bystander condition with IL-25/IL-33. Similarly, Granulocyte-macrophage colony-stimulating factor (GM-CSF) was secreted more efficiently with IL-1β/IL-23 and TCR stimulation than with the cytokines alone (Fig. 2K). Collectively, these results indicate that MP CD4^+^ T cells have functional heterogeneity when responding to specific sets of IL-1 family and STAT activators in the absence of their cognate antigen and suggest possible potent transcriptional regulators that control MP CD4^+^ T cell effector functions.

**Figure 2.**
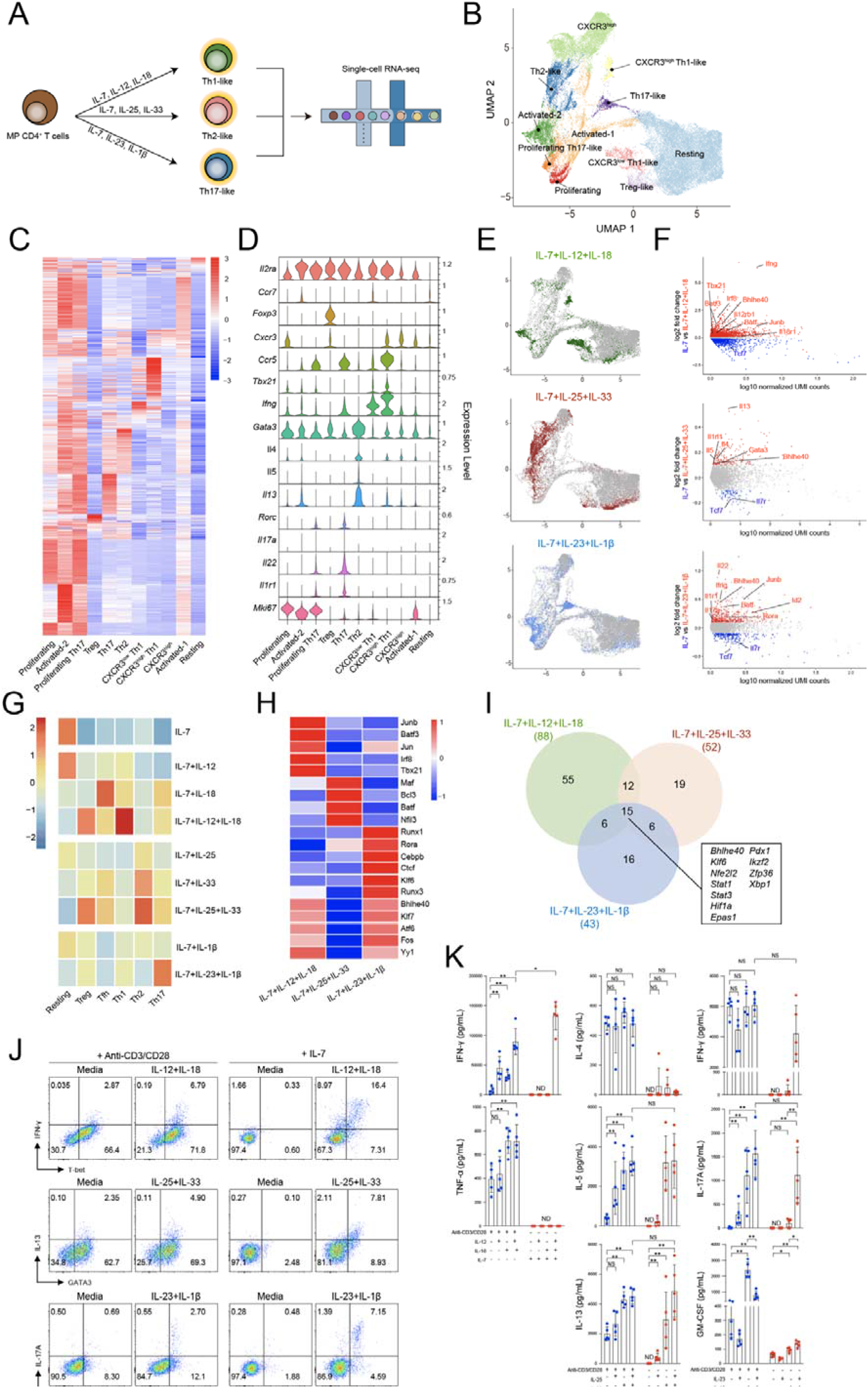
Phenotypic differences of MP CD4^+^ T cells after exposure to IL-1 family and STAT activating cytokines. (A) Overview of experimental design. (B) UMAP representation of MP CD4^+^ T cells stimulated with IL-12 and/or IL-18 and IL-33 and/or IL-25 and IL-1β and/or IL-23 in the presence of IL-7. (C) Heatmap and (D) violin plots of differentially expressed transcripts in cluster. (E) Individual cytokines conditions visualized with UMAP. (F) MA plots of differentially expressed genes comparing IL-7 versus IL-12/18 or IL-33/25 or IL-1β/23. (G) Heatmap representing gene expression of Resting-, Tfh-, Th1-, Th2-, Th17- and Treg-related gene signatures in each cytokine condition. (H) Heatmap representing transcription activity. (I) Venn diagram of transcriptional regulators predicted by the IPA. Numbers indicate the number of gene in each gate.(J) MP CD4^+^ T cells (TCRβ^+^CD1d tetramer^-^CD8^-^CD4^+^CD25^-^CD62L^low^CD44^high^) were cultured for 5 days with IL-12 and/or IL-18 and IL-33 and/or IL-25 and IL-1β and/or IL-23 in the presence of IL-7 or anti-CD3/anti-CD28. Representative flow cytometry plots showing the expression of effector lineage markers in each cytokine condition. (K) IFN-γ, TNF-α, IL-4, IL-5, IL-13, IL-17A, and GM-CSF were measured by ELISA (n=5). Data are presented as the mean ± S.D. P values were calculated using Mann-Whitney U-test (*p< 0.05, **p < 0.01, ***p < 0.001).

### Potential responder MP CD4^+^ T cells express distinct chemokine receptors upon IL-12/IL-18 and IL-1β/IL-23 stimulation

To determine which MP CD4^+^ T cells are potential responders to IL-1 family and STAT activating cytokines, we analyzed the subpopulation of expanding or responding clusters and performed trajectory analyses. Control MP CD4^+^ T cells cultured with IL-7 produced a significantly separate population of CXCR3^high^ cells. In IL-18/IL-12-conditioned MP CD4^+^ T cells the CXCR3^high^ cell population was reduced, and the IFN-γ- or IL-13^high^expressing Th1 and proliferating Th1 populations were greatly increased (Fig. 3A and 3B). Th1 signature genes, *Tbx21, Ifng, Cxcr3, Il2rb1, Il18r1*, and *CCR5* were highly expressed by IL-18/IL-12-conditioned MP CD4^+^ T cells (Fig. 3C). GO/KEGG analysis indicated that the related gene sets in those cells had increased including “Cellular response to interferon-gamma”, “Cytokine cytokine receptor interaction”, “Alzheimer’s disease”, and “Parkinson’s disease” (Fig. 3D). The IPA returned terminologies related to cytokines and inflammation such as “JAK/STAT signaling” and “neuroinflammation signaling pathway” and related to cytotoxic response including “Granzyme B signaling” (Fig. 3E). In a pseudo-time trajectory analysis, the MP CD4^+^ T cells formed a continuous progression that started in CXCR3^high^ cells and gradually progressed toward Fate 1, which expressed *Ifng, Stat5a, Bhlhe40, Batf3, Irf4* and *Irf8* (Fig. 3F and 3G). Similarly, a CCR6^high^ cluster was present in IL-7-conditioned MP CD4^+^ T cells, and its Th17-like cluster was specifically increased by IL-1β and IL-23 (Fig. 3H and 3I). Th17 signature genes, *Rorc, Ccr6, Il17a Il1r1, Il23r*, and proliferating marker *Mki67* were expressed by those cells (Fig. 3J). GO/KEGG analysis predicted that the related gene sets in IL-1β /IL-23-cultured MP CD4^+^ T cells had increased “Alzheimer’s disease,” “Parkinson’s disease,” and “Huntington’s disease” which are neurological diseases and also increased “cytokine signaling pathway” (Fig. 3K). The IPA more clearly explained the related pathways, “STAT3 pathway,” “leukocyte extravasation signaling,” “neuroinflammation signaling pathway,” “chemokine signaling,” and “Th17 activation pathway” (Fig. 3L). In the trajectory analysis, CCR6^high^ cells seemed to be the starting point, and then the cells gradually differentiated toward Fate 2 (Fig. 3M), which expresses pathogenic Th17-related genes such as *Rorc, Il17a, Csf2, Il22, Bhlhe40, Rora and Cebpb* with increased activities (Fig. 3N) and expression level (Fig. 3O) of related transcriptomes. These results collectively reveal that Th1-like and Th17-like MP CD4^+^ T cells expressing different chemokine receptors respond specifically to IL-12/IL-18 and IL-1β/IL-23 cytokines with the effector functions.

**Figure 3.**
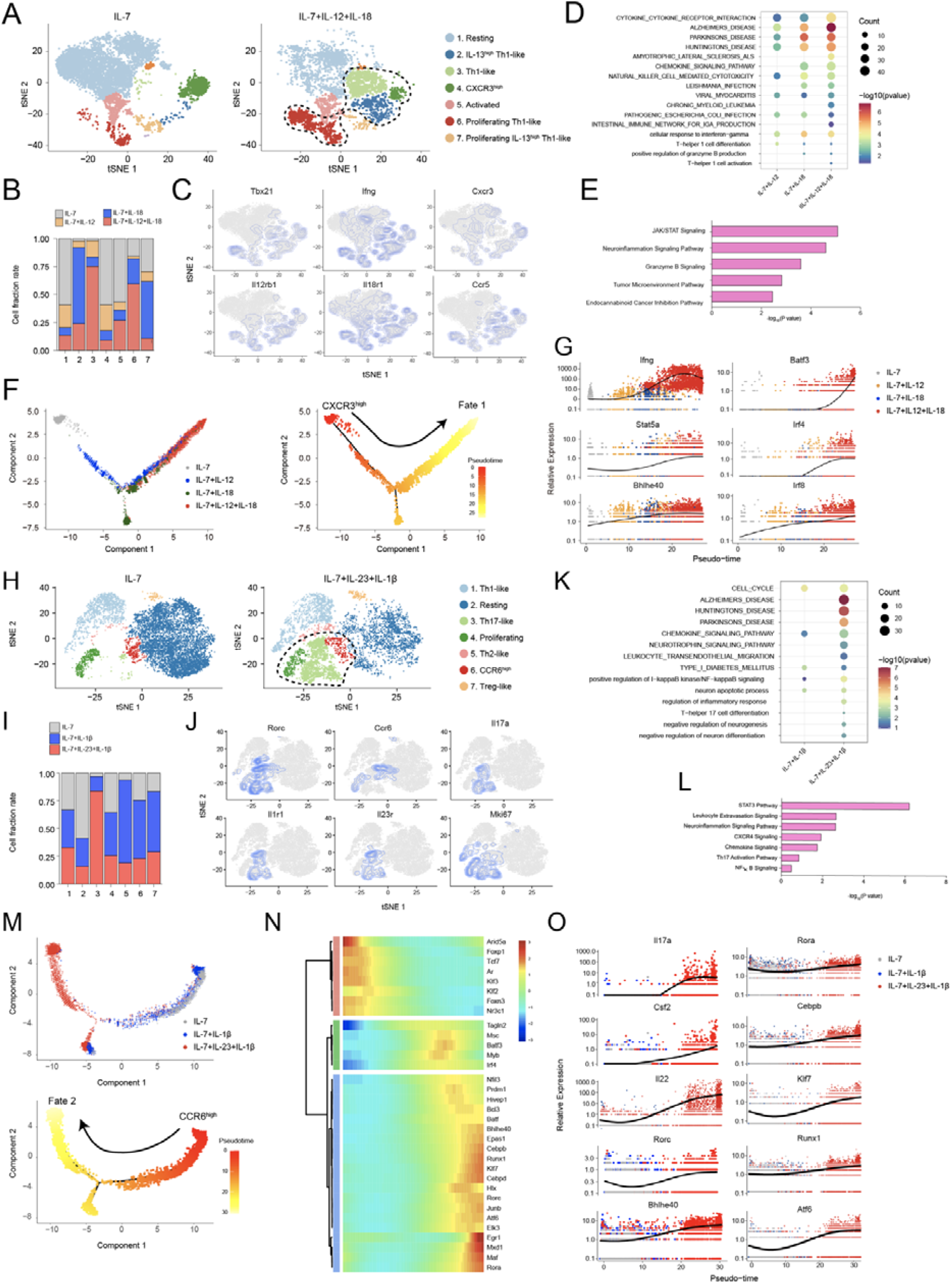
Potential responder MP CD4^+^ T cells express distinct chemokine receptors upon IL-12/IL-18 and IL-1β /IL-23 stimulation. (A) tSNE plots and (B) proportion of MP CD4^+^ T cells stimulated with IL-12 and IL-18 in the presence of IL-7 for 5 days. (C) Expression of selected Th1-related genes. (D) Selected KEGG/GO terms in cluster 2,3,4 and 6 and (E) Ingenuity pathway analysis (IPA) in cluster 3,4 and 6 of IL-12- and IL-18-responded MP CD4^+^ T cells. (F) Pseudo-time trajectory: each cell is colored by its pseudo-time value and (G) the expression level of the related genes. (H) tSNE plots and (I) proportion of MP CD4^+^ T cells stimulated with IL-1β and IL-23 in the presence of IL-7 for 5 days. (J) Expression of selected Th17-related genes. (K) Selected KEGG/GO terms and (L) IPA of IL-1β- and IL-23-stimulated MP CD4^+^ T cells (clusters 3, 4, 6). (M) Pseudo-times trajectory: each cell is colored by its pseudo-time value and (N) the transcription factor activity and (O) expression level of the related genes.

### CCR6^high^ memory phenotype CD4^+^ T cells are bystander-activated by IL-1β and IL-23 to become pathogenic Th17-like cells

Since scRNA-seq predicted that CCR6^high^ cells were the major cells responding to IL-1β and IL-23, we sorted splenic CCR6^high^ and CCR6^low^ MP CD4^+^ T cells (Fig. 4A and 4B) and then determined their proportions in steady-state SPF- and GF-housed mice (Fig. 4C). We found that splenic CCR6^high^ MP CD4^+^ T cells were independent of the gut (Fig. 4C). CCR6^high^ MP CD4^+^ T cells expressed RORγt more highly than CCR6^low^ cells and had comparable expression of T-bet (Fig. 4D and 4E). Steady-state CCR6^high^ MP CD4^+^ T cells, but not CCR6^low^, expressed IL-17A (Fig. 4F and 4G), suggesting that CCR6^high^ MP CD4^+^ T cells are Th17-like cells. Further stimulation by IL-1β and IL-23 induced IL-17A and GM-CSF expression (Fig. 4H and 4I), which confirms that CCR6^high^ MP CD4^+^ T cells produce pathogenic cytokines in the bystander manner. The amount of cytokine secreted, as determined by ELISA, consistently showed that CCR6^high^ MP CD4^+^ T cells, but not CCR6^low^, significantly produced IL-17A, GM-CSF, and IFN-γ and that IL-1β and IL-23 had important synergy for pathogenicity (Fig. 4J). In addition, IL-1β greatly expanded RORγt expression in CCR6^high^ MP CD4^+^ T cells but not CCR6^low^ cells, whereas IL-23 somewhat inhibited the proportion of Ki67-expressing cells, suggesting that IL-1β is important for the proliferation of CCR6^high^ MP CD4^+^ T cells (Fig. 4K). To further confirm the functions of IL-1β and IL-23 in MP CD4^+^ T cells, we performed bulk-RNA seq with bystander-activated MP CD4^+^ T cells exposed to IL-1β and IL-23. In the heatmap DEG analysis, IL-1β increased the expression of proliferation-related gene such as *Mki67* and *cdk2*, whereas IL-23 alone did not have any significant effects on gene expression (Fig. 4L). IL-23 and IL-1β together significantly induced the expression of pathogenic genes such as *Bhlhe40, Il1r1, Csf2, Ifng, Il22* and *Il17a*. In support, Gene set enrichment analysis (GSEA) pathway enrichment plot of IL-1β vs. IL-1β and IL-23 demonstrated that the enrichment score for cell proliferation was higher with IL-1β, and the score of the pathogenic Th17 signature with IL-1β and IL-23 was higher than in the control group (Fig. 4M). Through these results, we understand that steady-state CCR6^high^ MP CD4^+^ T cells, which show Th17-like characteristics, are the major bystander-activated cells responding to IL-1β and IL-23, which potentiate the cells’ pathogenic character.

**Figure 4.**
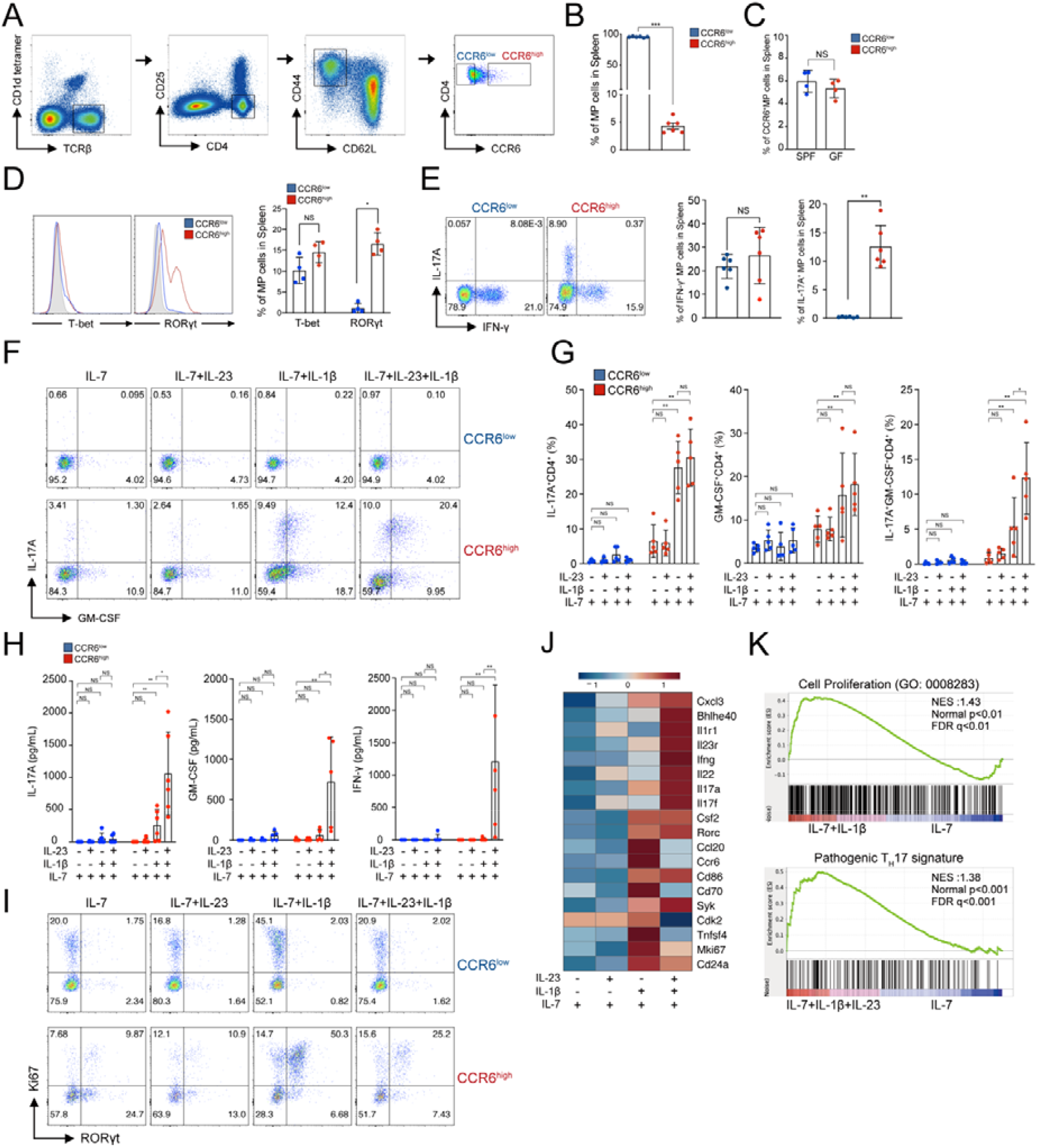
CCR6^high^ MP CD4^+^ T cells are bystander-activated by IL-1β and IL-23 to become pathogenic Th17-like cells. (A) Gating strategy of CCR6^high^ and CCR6^low^ MP CD4^+^ T cells from SPF mice. (B) The percentage of CCR6 expression in steady-state splenic MP CD4^+^ T cells (n=6). (C) The expression of CCR6 in SPF- vs. GF-MP CD4^+^ T cells (n=4). (dD) Transcription factor expression level of T-bet and RORγt in FACS-sorted CCR6^high^ and CCR6^low^ MP CD4^+^ T cells and (E) the proportion (n=4). (F) Representative flow cytometry plots showing the cytokine expression and (G) the average proportion of CCR6^high^ MP CD4^+^ T cells vs. CCR6^low^ MP CD4^+^ T cells (n = 6). CCR6^high^ and CCR6^low^ MP CD4^+^ T cells were stimulated with IL-1β and/or IL-23 in the presence of IL-7 for 5 days. (H) The representative proportion and (I) the average value showing the cytokine producing cells (n=5). (J) Concentration of IL-17A, GM-CSF, and IFN-γ were analyzed by ELISA (n = 5). (K) The proportion of RORγt^+^ and Ki67^+^ cells in CCR6^high^ and CCR6^low^ MP CD4^+^ T cells.(L) Heatmap of selected genes (M) Gene set enrichment analysis (GSEA) pathway enrichment plot related to “Cell Proliferation” and “Pathogenic T_H_17 signature” by bulk RNA-seq analysis. *q*, false discovery rate; NES, normalized enrichment score. Data are presented as the mean ± S.D. All data of p values were calculated using Mann-Whitney U-test (ND, not detected; NS, not significant; *p < 0.05, **p < 0.01, ***p < 0.001).

### CCR6^high^ MP CD4^+^ T cells are encephalitogenic and exacerbate autoimmune neuroinflammation

To reveal the importance of splenic MP CD4^+^ T cells during an autoimmune disease, we first induced EAE in 5-week-old mice, who have a lower proportion of MP CD4^+^ T cells than 10-week-old mice. In this mouse model, we compared the disease severity of control mice with that of those who received an additional adoptive transfer of Treg-deleted MP CD4^+^ T cells from 10-week-old Foxp3-GFP mice. EAE was rapidly induced and progressed by transferring additional MP CD4^+^ T cells from 10-week-old mice, suggesting that MP CD4^+^ T cells could be an important contributor to MOG_35-55_-induced EAE pathogenesis (Fig. S7A). In support, Rag^-/-^ mice that received MOG-TCR transgenic (2D2) naïve CD45.1^-^V_β_11+CD4^+^ T cells with Treg-deleted MP CD4^+^ T cells (CD45.1^+^CD4^+^) showed a more severe phenotype of EAE than the mice that received only 2D2 T cells (Fig. S7B), suggesting that MP CD4^+^ T cells contribute to the pathogenesis of EAE. Based on our previous results, we hypothesized that CCR6^high^ MP CD4^+^ T cells are the major pathogenic subpopulation contributing to EAE disease progression. To test that hypothesis, we transferred CCR6^high^ and CCR6^low^ MP CD4^+^ T cells (CD45.1^+^CD4^+^) and 2D2 naïve CD4^+^ T cells into Rag^-/-^ mice. The additional transfer of CCR6^high^ MP CD4^+^ T cells exacerbated EAE development compared with 2D2 transfer alone or the transfer of CCR6^low^ MP CD4^+^ T cells (Fig. 5A). Interestingly, the number of transferred MP CD4^+^ T cells that appeared in the spinal cord and brain tissue did not differ between conditions (Fig. 5B). However, CCR6^high^ MP CD4^+^ T cells produced significantly more cytokines, particularly IL-17A and GM-CSF, in the spinal cord and brain tissue than CCR6^low^ MP CD4^+^ T cells (Fig. 5C and D). Therefore, CCR6^high^ MP CD4^+^ T cells, along with antigen-specific T cells, contribute to the pathogenicity of autoimmune neuroinflammation by expressing pathogenic cytokines such as IL-17A and GM-CSF. To clarify the antigen-independent activation of CCR6^high^ MP CD4^+^ T cells in an EAE mouse model, we detected 2D2 TCR (V_α_3.2^+^ and V_β_11^+^), which are predominantly expressed in 2D2 naïve CD4^+^ T cells. Indeed, CCR6^high^ MP CD4^+^ T cells barely expressed V_α_3.2^+^ and V_β_11^+^ (Fig. 5E and F). In support, we confirmed that steady-state MP CD4^+^ T cells did not respond to the MOG_35-55_ antigen (Fig. 5G). Collectively, these results suggest that CCR6^high^ MP CD4^+^ T cells are encephalitogenic cells that exacerbate autoimmune neuroinflammation by producing IL-17A and GM-CSF, presumably in a bystander manner, along with antigen-specific T cells.

**Figure 5.**
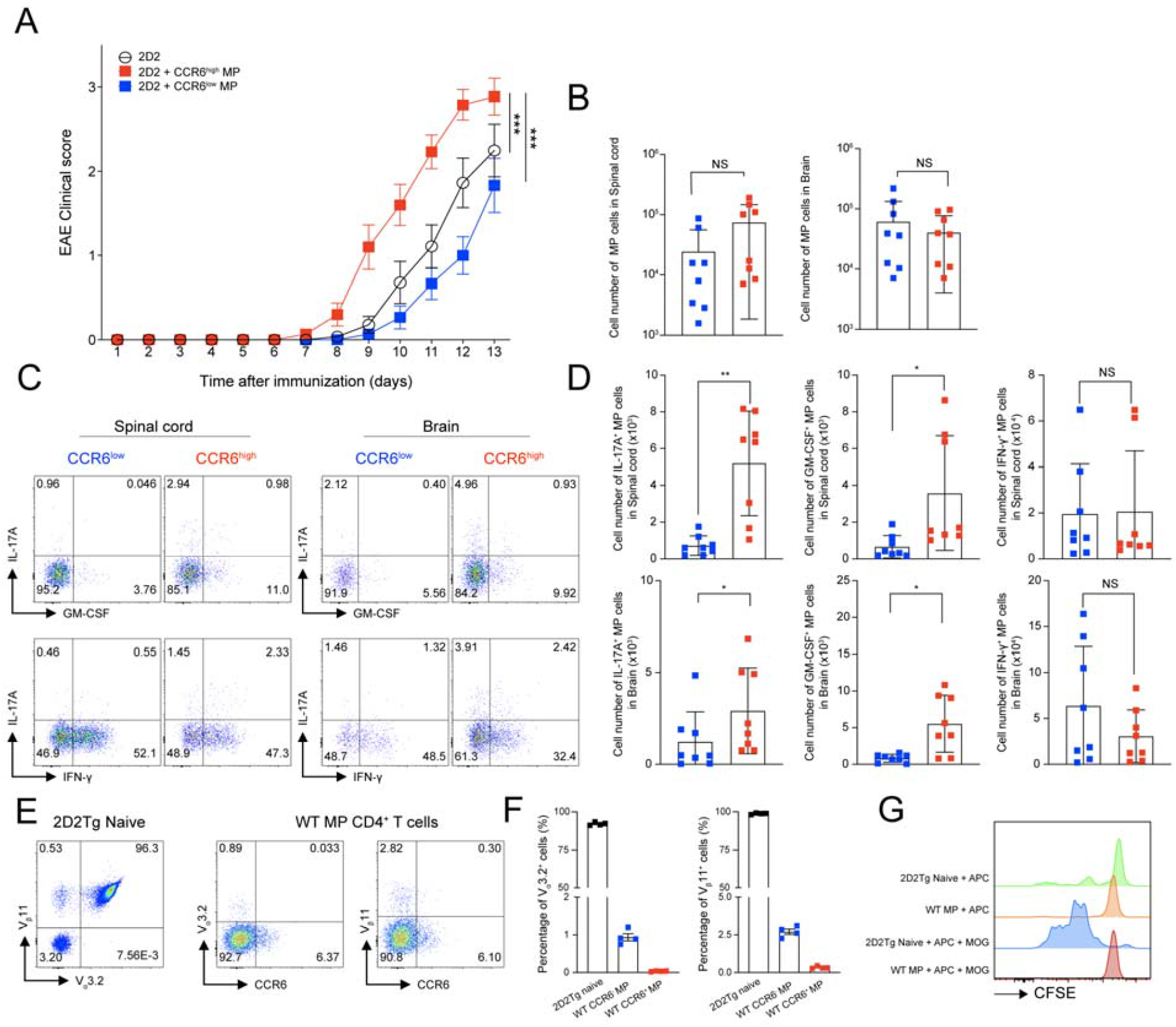
CCR6^high^ MP CD4^+^ T cells are encephalitogenic and exacerbate autoimmune neuroinflammation. (A) Naïve CD4^+^ T cells (5 × 10^4^) from 2D2 transgenic mice were adoptively transferred, with or without CCR6^high^ or CCR6^low^ MP CD4^+^ T cells (1 × 10^5^ CD45.1^+^TCRβ^+^CD1d tetramer^-^CD4^+^CD25^-^ CD62L^low^CD44^high^), into Rag^-/-^ mice who were immunized with MOG_35-55_ in CFA. EAE clinical score was monitored daily (n=15). (B) Absolute cell numbers of infiltrated CD45.1^+^MP CD4^+^ T cells in the spinal cord and brain (n=8). (C) The representative plots and (D) absolute cell number of IL-17A, GM-CSF, and IFN-γ producing cells were determined in MP CD4^+^ T cells from the spinal cords and brains on days 12–13 after immunization (n=8). (E) The representative dot plots and (F) the average value showing the percentage of 2D2 TCR (V_α_3.2^+^ and V_β_11^+^) in spleens from 2D2 transgenic mice and C57BL/6 wild type mice (n=4). (G) FACS-sorted 2D2 naïve CD4^+^ T cells and MP CD4^+^ T cells were cultured with CD11c^+^ dendritic cells (MHC-II^+^CD11c^+^) with or without MOG_35-55_ peptide (50 μg/ml) for 3 days. CFSE levels were measured by flow cytometry. Data are presented as the mean ± S.E.M in A and the mean ± S.D in B,D,F. *P* values were calculated using two-way ANOVA or Mann-Whitney U-test (NS, not significant; *p < 0.05, **p < 0.01, ***p< 0.001).

### Bhlhe40 confers pathogenic functions of bystander CCR6^high^ MP CD4^+^ T cells by IL-1 and IL-23

Among the candidate genes involved in bystander activation of CCR6^high^ MP CD4^+^ T cells, we identified that Bhlhe40/GM-CSF axis could potentially give rise to the pathogenic function of CCR6^high^ MP CD4^+^ T cells induced by IL-1β and IL-23 without TCR stimulation (Fig. 6A). By using Bhlhe40^GFP^ mice, we found that CCR6^high^ MP CD4^+^ T cells activated by IL-1β and IL-23 showed significantly increased level of Bhlhe40 (Fig. 6B). Interestingly, Bhlhe40^GFP^ positive T cells majorly produced effector cytokines including IL-17A and GM-CSF compared to Bhlhe40^GFP^ negative T cells (Fig. 6C). In support, Bhlhe40^-/-^ CCR6^high^ MP CD4^+^ T cells showed markedly reduced IL-17A and GM-CSF production compared to WT (Fig. 6D and E), suggesting that Bhlhe40 is an important transcriptional regulator for the pathogenic functions of bystander CCR6^high^ MP CD4^+^ T cells. To confirm the in vivo relevance, we transferred WT CCR6^high^ or Bhlhe40^-/-^ CCR6^high^ MP CD4^+^ T cells along with 2D2 naïve CD4^+^ T cells into Rag^-/-^ mice. Bhlhe40^-/-^ CCR6^high^ MP CD4^+^ T cells showed abrogated functions for exacerbating EAE disease compared by WT CCR6^high^ MP CD4^+^ T cells (Fig. 6F). Interestingly, the number of Bhlhe40^-/-^ CCR6^high^ MP CD4^+^ T cells expressing IL-17A and GM-CSF in the spinal cord and brain tissue was significantly reduced compared to WT CCR6^high^ MP CD4^+^ T cells (Fig. 6G and H) without significant difference of MOG-specific 2D2 T cells (data not shown). In addition, transfer of GM-CSF-deficient CCR6^high^ MP CD4^+^ T cells showed abrogated function of contributing EAE pathogenesis (Fig. 6I), suggesting that disease aggravation by bystander CCR6^high^ MP CD4^+^ T cells is committed by IL-1β and IL-23 via Bhlhe40/GM-CSF axis. Collectively, these results indicate that Bhlhe40 confers pathogenic functions of bystander CCR6^high^ MP CD4^+^ T cells with GM-CSF production.

**Figure 6.**
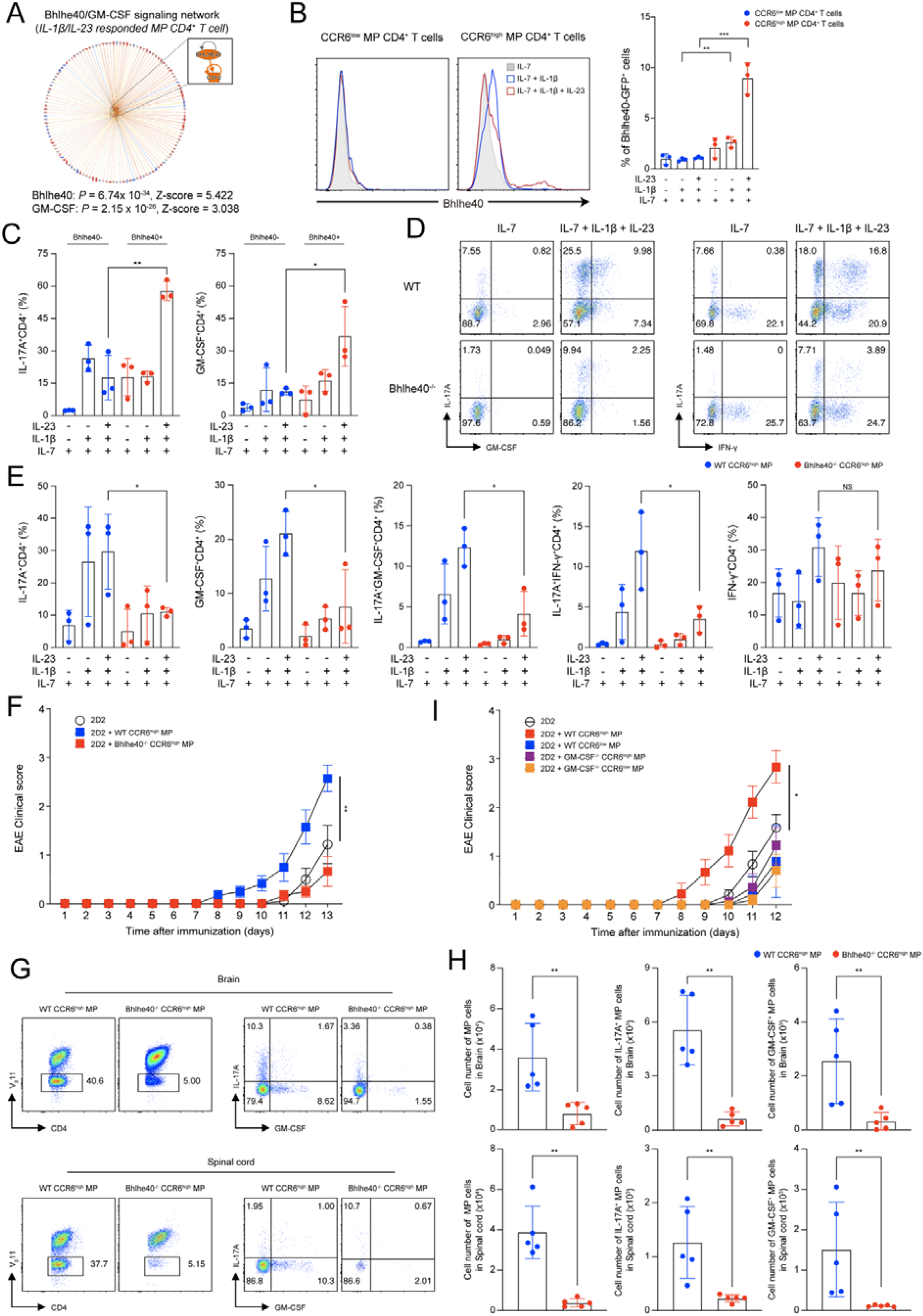
Bhlhe40 confers pathogenic functions of bystander CCR6^high^ MP CD4^+^ T cells by IL-1 and IL-23. (A) Predicted upstream network on IL-1β and IL-23 responded MP CD4^+^ T cells by IPA. (B) The representative histogram and average value showing the percentage of Bhlhe40 level in CCR6^high^ and CCR6^low^ MP CD4^+^ T cells induced by IL-1β/IL-23 stimulation without TCR engagement (n=3). (C) The representative percentage of IL-17A and GM-CSF expression compared to Bhlhe40^GFP^ positive and negative cells in CCR6^high^ MP CD4^+^ T cells induced by IL-1β /IL-23 without TCR engagement for 5 days (n=3). (D) Representative flow cytometry plots showing the cytokine expression and (E) average proportion of WT CCR6^high^ MP CD4^+^ T cells vs. Bhlhe40^-/-^ CCR6^high^ MP CD4^+^ T cells (n = 3). (F) Naïve CD4^+^ T cells (5 × 10^4^) from 2D2 transgenic mice were adoptively transferred, with or without WT CCR6^high^ or Bhlhe40^-/-^ CCR6^high^ MP CD4^+^ T cells (1 × 10^5^, TCRβ^+^CD1d tetramer^-^CD4^+^CD25^-^CD62L^low^CD44^high^), into Rag^-/-^ mice who were immunized with MOG_35-55_ in CFA. EAE clinical score was monitored daily (n=5). (G) Representative flow cytometry plots and (H) absolute cell numbers of infiltrated 2D2 TCR (V_β_11^+^)^-^ MP CD4^+^ T cells (gating from CD45^+^CD4^+^) and IL-17A, GM-CSF, and IFN-γ producing cells from the spinal cords and brain on days 13 after immunization (n=5). (I) Naïve CD4^+^ T cells (5 × 10^4^) from 2D2 transgenic mice were adoptively transferred, with or without WT CCR6^high^ or WT CCR6^low^ or GM-CSF^-/-^ CCR6^high^ or GM-CSF^-/-^ CCR6^low^ MP CD4^+^ T cells (1 × 10^5^, CD45.1^+^TCRβ^+^CD1d tetramer^-^CD4^+^CD25^-^CD62L^low^CD44^high^), into Rag^-/-^ mice who were immunized with MOG_35-55_ in CFA. EAE clinical score was monitored daily (n=9). Data are presented as the mean ± S.E.M in f,i and the mean ± S.D in b,c,d,e,g,h values were calculated using two-way ANOVA or Mann-Whitney U-test (NS, not significant; *p < 0.05, **p < 0.01, ***p < 0.001).

## Discussion

In this study, we intensively validated the characteristics of MP conventional CD4^+^ T cells using scRNA-seq to unveil their effector functions during autoimmune disease. We found distinct effector-like subpopulations in steady-state splenic MP CD4^+^ T cells that are independent of the microbiome and food antigens. Moreover, MP CD4^+^ T cells can be bystander-activated toward Th1-, Th2-, and Th17-like effector cells by the IL-1 family and STAT activating cytokines. In particular, chemokine receptor–expressing cells are defined as potential responder cells, and we focused on CCR6^high^ MP CD4^+^ T cells to further validate their functions in responding to IL-1β/IL-23. We demonstrated that encephalitogenic CCR6^high^ MP conventional CD4^+^ T cells are bystanders that exacerbate EAE disease progression, along with antigen-specific T cells. Furthermore, we suggested Bhlhe40 as a pivotal transcriptional regulator that governs GM-CSF production by CCR6^high^ MP CD4^+^ T cells, exacerbating EAE development. Overall, our results reveal the immunological function of antigen-unrelated MP CD4^+^ T cells in autoimmune disease.

According to previous reports, CD44^high^ MP CD4^+^ and CD8^+^ conventional T cells exist spontaneously in mouse spleens, independently of germ and food antigens, and also in humans (Byrne *et al*., 1994; Haluszczak *et al*., 2009; Kawabe *et al*., 2017; Szabolcs *et al*., 2003). In support, we and others confirmed that MP CD4^+^ T cells are found in mice even 1-2 days after birth and that they increase significantly with age in both the spleen and other tissues (Kawabe *et al*., 2017). However, the generation of MP CD8^+^ T cells remains controversial. Some studies have reported that MP CD8^+^ T cells are derived from naïve T cells in lymphopenic conditions (Haluszczak *et al*., 2009; Hamilton *et al*., 2006; Seddon *et al*, 2000; White *et al*., 2016), whereas other groups have shown that MP CD8^+^ T cell generation is a TCR-related process triggered by the recognition of self-antigens in the thymus (Miller *et al*, 2020). Along the same lines, MP CD4^+^ T cells have recently been reported to differentiate from naïve CD4^+^ T cells in the periphery in both neonates and adult mice in a thymus-independent fashion (Kawabe *et al*., 2017). However, it has been recently reported that there is a proportion of memory CD4^+^ T cells in human thymus by scRNA-seq (Park *et al*, 2020), suggesting the potential generation of natural MP T cells. In support, we observed a significant proportion and number of CD44^high^ MP CD4^+^ T cells in the mouse thymus that increased with age. Collectively, therefore, these studies do not rule out the possibility that MP CD4^+^ T cells might be generated both from naïve T cells and directly by the thymus. Furthermore, we found that the TCR diversity of steady-state MP CD4^+^ T cells was slightly reduced in GF mice, revealing that the gut microbiome potentially contributes to certain minor TCR clonotypes of peripheral MP CD4^+^ T cells. In the mLNs, Th17-like MP CD4^+^ T cells were significantly reduced in GF-housed mice, suggesting the importance of the microbiome to gut Th17 cells. Therefore, peripheral self-antigens and the gut microbiome might both contribute to TCR diversity and the proportion of MP CD4^+^ T cells.

The heterogeneous characteristics of MP conventional CD4^+^ T cells were investigated using single cell transcriptomics, which revealed clearly distinct Th1-, Th17-, Tfh-, and Treg-like subpopulations. In support, recent studies revealed the potential heterogeneity of murine MP CD4^+^ T cells (ElTanbouly *et al*, 2020; Kawabe *et al*, 2020) and further demonstrated that a CXCR3^+^T-bet^+^ Th1-like MP population was differentiated from naïve CD4^+^ T cells by DC1-derived IL-12 and further activated by CD40–CD40L interactions between DC1 and CD4^+^ T cells (Kawabe *et al*., 2020). Of note, MP CD4^+^ and CD8^+^ T cells rapidly produce IFN-γ in response to IL-12 and IL-18 (Guo *et al*, 2009; Kawabe *et al*., 2017; Sosinowski *et al*., 2013; White *et al*., 2017). Given those observations, we hypothesized that CXCR3^high^ MP CD4^+^ T cells are major responders to IL-12 and IL-18 cytokine stimulation. As expected, we confirmed that CXCR3^high^ MP CD4^+^ T cells, in high correlation with CCR5 expression, can produce IFN-γ and T-bet in response to IL-12 and IL-18, suggesting a potential innate-like function of CXCR3^high^ MP CD4^+^ T cells. Interestingly, without TCR stimulation, cytokine-mediated bystander activation could not induce TNF-α from MP CD4^+^ T cells, suggesting that TCR and cytokine signaling produce distinct effector functions. We also used IL-33 and IL-25 stimulation of MP CD4^+^ T cells to see whether they produced Th2 cytokines such as IL-13 and IL-5. Interestingly, the MP CD4^+^ T cells produced IL-4 with TCR stimulation, suggesting that the effector function of MP CD4^+^ T cells bystander-activated by IL-33 and IL-25 correlates with innate lymphoid cell type 2 (ILC2). A previous report consistently showed that *in vitro*–generated memory Th2 cells mainly produce IL-13, rather than IL-4, upon exposure to IL-33 and IL-7 without a cognate antigen. In addition, like ILC2, tissue-resident memory Th2 cells were activated by IL-33 and showed a protective function against helminth infection in an antigen-independent manner (Guo *et al*, 2015). Although we did not observe a significant Th2-like MP subpopulation in the steady state, we confirmed that Th2-like populations are induced by IL-33 and IL-25, suggesting that heterogeneous IL-33R- or IL-25R-expressing cells might be responding to those cytokines.

In our previous study, we demonstrated that and IL-23 can synergistically potentiate the pathogenicity of memory CD4^+^ T cells *in vitro* (Lee *et al*., 2019) and that non-myelin-specific CD4^+^ T cells can infiltrate the CNS with MOG antigen–specific T cells, which significantly contribute to EAE disease progression (Jones *et al*., 2003; Lee *et al*., 2019). In rheumatoid arthritis patients, T cells that infiltrate the synovial fluid mainly express the CD45RO^+^ memory marker and specifically respond to epitopes of Epstein-Barr virus and cytomegalovirus (Kobayashi *et al*., 2004; Pacheco *et al*, 2019; Scotet *et al*, 1999; Tan *et al*., 2000). In type 1 diabetes, infection with rotavirus or coxsackie virus is reported to be involved in accelerated diabetes onset through Toll-like receptor (TLR) signaling without pancreatic infection (Horwitz *et al*, 1998; Pane *et al*, 2014), and influenza A virus is linked to diabetes in human patients (Nenna *et al*, 2011; Pane & Coulson, 2015). Collectively, those studies suggest that antigen-non-related CD4^+^ T cells can contribute to disease onset or progression with antigen-specific T cells in various autoimmune diseases. We have identified here that CCR6^high^ MP CD4^+^ T cells are the major subpopulation of MP CD4^+^ T cells that respond to IL-1β and IL-23 by expanding and inducing pathogenic Th17 characteristics. In an adoptive transfer model of EAE, CCR6^high^ MP CD4^+^ T cells transferred with MOG-specific T cells induced more severe EAE than CCR6^low^ cells, with increased production of IL-17 and GM-CSF in the CNS. We further confirmed that MP CD4^+^ T cells do not respond to MOG_33-55_ antigen, indicating the bystander role of CCR6^high^ MP CD4^+^ T cells in autoimmune neuroinflammation. Because MOG-specific T cells produce effector cytokines such as IL-17 and GM-CSF (data not shown), further studies should be done to reveal the distinct mechanism between antigen-specific T cells and bystander-activated T cells in autoimmune disease pathogenesis.

Analyzing the single cell transcriptomics of IL-1β/IL-23 responding MP CD4^+^ T cells, we identified that Bhlhe40 could be a potential transcriptional regulator inducing GM-CSF in CCR6^high^ MP CD4^+^ T cells. Bhlhe40 has been reported to play pivotal roles in T cells. Bhlhe40-deficient naïve CD4^+^ T cells show limited response to TCR stimulation (Martinez-Llordella *et al*, 2013). Moreover, Bhlhe40 seem to be required for Th1 and Th17 effector cytokine production including IL-17A, GM-CSF and IFN-γ in the context of autoimmune disease, GVHD, and Toxoplasma gondii infection model (Lin *et al*., 2016; Lin *et al*, 2014; Piper *et al*, 2020; Yu *et al*, 2018). In addition, a binding site for Bhlhe40 was identified the correlation of mouse *Csf2* locus, which encodes GM-CSF, (Jarjour *et al*, 2020; Lin *et al*., 2014) and Bhlhe40 expression positively correlate with GM-CSF expression in human PBMC (Emming *et al*, 2020). We demonstrate that steady-state CCR6^high^ MP CD4^+^ T cells have Th17-like character with expressing RORγt. They up-regulated Bhlhe40 in response to IL-1β and IL-23 in the absence of TCR engagement for IL-17 and GM-CSF production. Moreover, transferring these cells in vivo, we found that Bhlhe40 is required in bystander CCR6^high^ MP CD4^+^ T cells for the expression of GM-CSF contributing to the pathogenesis of EAE development. In support, a previous study reported the majority of Bhlhe40-expressing pathogenic T cells in active EAE are non-MOG-specific (Lin *et al*., 2016; Lin *et al*., 2014). Therefore, Bhlhe40 can be a pivotal transcriptional regulator for both antigen-specific and bystander MP T cells in the context of CNS inflammation and targeting of Bhlhe40 in CD4^+^ T cells may serve as a potential novel treatment strategy to control autoimmune diseases.

Innate T cells such as natural killer T (NKT), mucosal-associated invariant T (MAIT), and γδ T cells have limited TCR gene usage compared to conventional T cells, which recognize complexes of non-peptide antigens such as glycolipids, phosphoantigens, and vitamin B metabolites, respectively (Godfrey *et al*, 2015; Lee *et al*, 2020; Pellicci *et al*, 2020). These innate T cells are derived from the thymus, which can evoke robust cytokine production. Previously, innate lymphocytes, such as NKT17, γδT17 cell, and ILC3 subsets have been defined to commonly express IL-17 and RORγt (Lee *et al*, 2013b; Sutton *et al*, 2009; Walker *et al*, 2013). Furthermore, a recent study reported a novel subset of αβ-γδ co-expressing T cells which recognize MHC-restricted peptide antigens and produce effector cytokine IL-17A, GM-CSF, and IFN-γ by IL-1β and IL-23 stimulation (Edwards *et al*, 2020). In this context, we carefully eliminated the possibility of contamination of innate T cells by sorting conventional MP CD4^+^ T cells using NKT (CD1d tetramer), γδ T, αβ-γδ T, ILC, and MAIT (TCRβ^+^CD8^-^CD4^+^CD25^-^CD44^high^CD62L^low^) exclusion gates. We confirmed that PLZF/CD1d tetramer negative steady-state MP CD4^+^ T cells still exists as a heterogenous population containing CCR6^high^RORγt^+^IL-17A^+^ cells, which is majorly bystander-activated by IL-1β and IL-23. Therefore, collectively, we provide compelling evidence that conventional CD4^+^ T cells distinguished from previously known innate T cells exist, which have an innate-like features and contribute to autoimmune neuroinflammation.

Memory T cells are antigen-experienced cells that respond to their cognate antigens more rapidly upon re-exposure. Therefore, we speculate that MP CD4^+^ T cells would have antigen-specific memory responses, and we wonder what the antigen is. Previous studies showed that antigen-specific cells called virtual memory cells exist without antigen exposure and nonetheless perform the memory response. For example, *vaccinia* virus B8R- and HSV-specific MP CD8^+^ T cells in unimmunized mice can react to those specific antigens (Haluszczak *et al*., 2009; Hamilton *et al*., 2006). Virtual memory CD8^+^ T cells have a CD44^high^CD122^+^CD49d^low^ phenotype which requires a transcription factor *Eomes* that differs from that of antigen-induced memory cells (Sosinowski *et al*., 2013). In addition, virus-specific MP CD4^+^ T cells are abundant in unexposed adults, presumably through cross-reactivity, so that they have the typical properties of memory T cells (Su & Davis, 2013; Su *et al*, 2013). However, virtual memory CD8^+^ T cells are also known to play a protective role in infection using an antigen-non-specific manner (White *et al*., 2016). Therefore, MP CD4^+^ T cells derived from steady-state mice might have cross-reactivity to certain pathogens even though they did not respond to the MOG_35-55_ antigen *in vitro*, and we speculate that they have both antigen-specific and bystander functions.

Antigen-specific T cells are fundamentally important in triggering autoimmune inflammation; however, antigen-non-related MP CD4^+^ T cells also contribute to pathogenic inflammation in a bystander manner. Collectively, our studies of the role that MP CD4^+^ T cells play in neuroinflammatory disease shed light on the innate-like function of adaptive immune cells to understand disease pathogenesis and reveal a novel drug development strategy to modulate autoimmune diseases.

## Experimental and Methods

### Mice

C57BL/6J mice were purchased from DBL (Chungcheongbuk-do, Korea), and Rag^−/−^, GM-CSF^-/-^ and 2D2 TCR-transgenic mice were purchased from Jackson Laboratory (Bar Harbor, ME, USA). CD45.1^+^, Foxp3-GFP mice were provided by Jeehee Youn (Hanyang University). GF and AF mice were purchased from the animal facility of POSTECH Biotech Center (Pohang, Korea). Bhlhe40^-/-^ and Bhlhe40^GFP^ mice were provided by Brian T. Edelson (Washington University). The mice were housed and bred in a specific pathogen–free animal facility at Hanyang University under controlled conditions with a constant temperature (21 ± 1 °C) and humidity (50 ± 5%) and a 12 h light/dark cycle with regular chow and autoclaved water.

### Cell isolation and differentiation

MP (TCRβ^+^CD4^+^CD1d tetramer^−^CD25^−^CD62L^low^CD44^high^) CD4^+^ T cells from the spleens of 8 to 12-week-old mice were isolated using a FACS Aria cell sorter II (BD Biosciences, Franklin Lakes, NJ, USA). FACS-sorted MP CD4^+^ T cells were stimulated with IL-1β (20 ng/mL, R&D Systems, Minneapolis, MN, USA), IL-23 (20 ng/mL, R&D Systems), IL-12 (20 ng/mL, Peprotech, Rocky Hill, NJ, USA), IL-18 (20 ng/mL, R&D Systems), IL-33 (20 ng/mL, R&D Systems), IL-25 (20 ng/mL, R&D Systems), IL-7 (10 ng/mL, Peprotech, Rocky Hill, NJ, USA), or plate-bound anti-CD3/anti-CD28 (2 μg/mL, BD Biosciences) for 5 days at 37 °C in an incubator.

### Flow cytometry

Cell surface staining was performed using the following monoclonal antibodies: anti-CD4 (RM4-5; eBioscience, San Diego, CA, USA, dilution 1:500), anti-CD25 (PC61.5; eBioscience, dilution 1:500), anti-CD44 (IM7; BioLegend, San Diego, CA, USA, dilution 1:500), anti-CD62L (MEL-14; BioLegend, dilution 1:500), anti-CD45 (30-F11, BioLegend, dilution 1:500), anti-CD45.1 (A20; eBioscience, dilution 1:500), anti-TCRβ (H57-597; eBioscience, dilution 1:500), anti-CCR6 (29-2L17; BD, dilution 1:100), anti-CXCR3 (CXCR3-173; BD, dilution 1:100), anti-V_α_3.2 (RR3-16; BioLegend, dilution 1:500), anti-V_β_11 (RR3-15; BioLegend, dilution 1:500), and PE-conjugated CD1d tetramer (PBS57; NIH, dilution 1:1000). Biotinylated PBS57 loaded and unloaded CD1d monomers were provided by the US National Institutes of Health Tetramer Core facility. For intracellular staining, the cells were stimulated with a cell stimulation cocktail (00-4975-03; eBioscience) for 4 h at 37 °C, and then surface markers were stained. After staining of the surface markers, the cells were fixed and permeabilized in Cytofix/Cytoperm (554714; BD Bioscience) or FOXP3/Transcription factor staining buffer set (00-5523-00; eBioscience) for 30 min at 4 °C or RT. Intracellular staining was performed using the following monoclonal antibodies: anti-IL-17A (eBio17B7; eBioscience, dilution 1:200), anti-IFN-γ (XMG1.2; eBioscience, dilution 1:400), anti-GM-CSF (MP1-22E9; BD Biosciences, dilution 1:200), anti-Ki67 (SolA15; eBioscience, dilution 1:500), anti-IL-4 (11B11; BioLegend, dilution 1:200), IL-13 (eBio13A; eBioscience, dilution 1:200), anti-TNF-α (MP6-XT22; BioLegend, dilution 1:500), anti-IL-5 (TRFK5; BioLegend, dilution 1:200), anti-IL-1R1 (35F5; BD Biosciences, dilution 1:100), anti-RORγt (Q31-378; BD Biosciences, dilution 1:100), anti-T-bet (4B10; BioLegend, dilution 1:50), anti-GATA3 (L50-823; BD Bioscience, dilution 3 μl per well) and IgG1 (R2-34; BD Biosciences). Stained cells were analyzed by flow cytometry (FACS Canto II, BD Bioscience), and data were analyzed using FlowJo software version 10.8.0 (Tree Star, Ashland, OR, USA).

### Experimental autoimmune encephalomyelitis

In EAE model, FACS-sorted MP CD4^+^ T cells (TCRβ^+^CD4^+^CD1d tetramer^-^Foxp3^-^CD62L^low^CD44^high^, 5 × 10^5^) from Foxp3-GFP mice were adoptively transferred to 5-week-old female C57BL/6 mice. After transfer, the mice were immunized with 200 μg of MOG_35-55_ peptide in complete Freund’s adjuvant (Chondrex, Inc., USA). At 0 and 48 h after immunization, the mice were intraperitoneally treated with 500 ng of pertussis toxin (List Biological Laboratories, Inc., Campbell, CA, USA). The animals were scored daily for clinical disease. In another EAE model by adoptive transfer, naive (CD4^+^V_β_11^+^CD25^-^CD62L^high^CD44^low^) CD45.1^−^ T cells (1-5 × 10^4^) from 2D2 TCR-transgenic mice were transferred into Rag^-/-^ mice with or without WT CCR6^high^ or WT CCR6^low^ or Bhlhe40^-/-^ CCR6^high^ or GM-CSF^-/-^ CCR6^high^ or GM-CSF^-/-^ CCR6^low^ MP CD4^+^ T cells (1.0 × 10^5^). Before transfer, CCR6^high^ or CCR6^low^ MP CD4^+^ T cells were primed *in vitro* with IL-7 (10 ng/mL) and IL-1β (20 ng/mL) for 5–7 days. After transfer, the mice were immunized with 100 μg of MOG_35−55_ peptide in complete Freund’s adjuvant (Chondrex, Inc., USA). At 0 and 48 h after immunization, the mice were intraperitoneally treated with 200 ng of pertussis toxin (List Biological Laboratories, Inc., Campbell, CA, USA). The animals were scored daily for clinical disease as follows (Stromnes & Goverman, 2006): partially limp tail, 0.5; completely limp tail, 1; limp tail and waddling gait, 1.5; paralysis of one hind limb, 2; paralysis of one hind limb and partial paralysis of the other hind limb, 2.5; paralysis of both hind limbs, 3; ascending paralysis, 3.5; paralysis of trunk, 4; moribund, 4.5; dead, 5. On day 12 or 13, the mice were sacrificed and perfused with Phosphate buffered saline (PBS). To isolate lymphocytes, the spinal cord and brain were digested with 1 mg/mL of collagenase D (11 088 866 001; Sigma-Aldrich) and DNase I (10 104 159 001; Sigma-Aldrich) and incubated at 80 RPM on a shaker for 35 min. After enzyme digestion, lymphocytes were isolated by Percoll (GE Healthcare, Little Chalfont, UK) density-gradient centrifugation.

### T cell proliferation and cytokine assays

For antigen-specific proliferation, 2D2 naïve (CD4^+^V_β_11^+^CD25^-^CD62L^high^CD44^low^) and MP CD4^+^ T cells (CD4^+^CD25^-^CD62L^low^CD44^high^) and CD11c^+^ dendritic cells (MHC-II^+^CD11c^+^) (APC) from the spleens of 2D2 TCR-transgenic and C57BL/6 mice were isolated using a FACS Aria cell sorter II. Before co-culture, naïve and MP CD4^+^ T cells were stained with 1.25 μM carboxyfluorescein succinimidyl ester (CFSE) (Invitrogen, Carlsbad, CA) for 7 min at room temperature. After incubation, 10% Fetal bovine serum (FBS) was added, and the incubation was continued on ice for 3 min. 2D2 Naïve and MP CD4^+^ T cells (1 × 10^5^ cells/well) were then washed with PBS and co-cultured with CD11c^+^ dendritic cells (5 × 10^4^) with or without 50 μg/mL of MOG_35-55_ peptide for 72 h.

### Enzyme-linked immunosorbent assay (ELISA)

Samples were measured using IL-17A ELISA (432501; BioLegend), IL-13 ELISA (88-7137-88; Thermo Fisher), IL-4 ELISA (431104; BioLegend), IL-5 ELISA (431204; BioLegend), IFN-γ ELISA (430801;BioLegend), TNF-α ELISA (430904; BioLegend), and GM-CSF ELISA (432201; BioLegend) kits according to the manufacturers’ instructions. Briefly, microwell plates (Corning Costar; 9018) were coated with capture antibodies overnight at 4 °C and blocked with ELISA diluent 1X for 1 h at room temperature. Samples and two-fold serial diluted standards were incubated at room temperature for 2 h, and then detection antibodies and streptavidin-HRP were added. Then, 1X TMB solution and a stop solution (2 N H_2_SO_4_) was loaded. The optical density was analyzed at 450 nm. Between all procedures, plates were washed at least 3 times with wash buffer (1X PBS, 0.05% Tween-20).

### Statistical analysis

All data were analyzed in non-parametric analyses using the Mann-Whitney test or two-way ANOVA of variance in Prism version 8.0 (GraphPad Software, San Diego, CA). Data are presented as the mean ± S.D. or mean ± S.E.M. For all data, significance was defined as *p* ≤ 0.05. Sample size and statistical information is provided in each figure legend.

### Single-cell RNA sequencing sample preparation

MP (TCR β^+^ CD4^+^ CD25^−^ CD62L ^low^ CD44 ^high^) CD4^+^ T cells from the spleens of 10-week-old mice were isolated using a FACS Aria cell sorter II (BD Biosciences, Franklin Lakes, NJ, USA). FACS-sorted MP CD4^+^ T cells were stimulated with IL-1β (20 ng/mL, R&D Systems, Minneapolis, MN, USA), IL-23 (20 ng/mL, R&D Systems), IL-12 (20 ng/mL, Peprotech, Rocky Hill, NJ, USA), IL-18 (20 ng/mL, R&D Systems), IL-33 (20 ng/mL, R&D Systems), IL-25 (20 ng/mL, R&D Systems), or IL-7 (10 ng/mL, Peprotech, Rocky Hill, NJ, USA) for 5 days at 37 °C in an incubator.

### Single-cell RNA- and TCR-seq data processing

Raw sequencing data were processed using a well-developed program, Cellranger (10X genomics), for quality control (QC) and gene expression estimation. Low-quality cells were filtered using 4 QC metrics: the percentage of mitochondrial gene unique molecular identifier (UMI) counts, the number of expressed genes (UMI count > 0), the sum of UMI counts per cell, and the doublet score estimated using Doublet Finder [10.1016/j.cels.2019.03.003]. The outlier cells from each QC metric were excluded from further analyses. To collect pure TCRα/β^+^ CD4^+^ T cells, cells expressing PLZF (*Zbtb16*) and *Cd74*^+^-cells in which at least 1 UMI was counted were removed. All TCRα/β^+^ CD4^+^T cells were integrated by cytokine treatment (set 1: IL-7, IL-7+ IL-1β, and IL-7+ IL-1β +IL-23; set 2: IL7, IL7+IL-25, IL-7+IL-33, and IL-7+IL-25+IL-33; set 3: IL-7, IL-7+IL-12, IL-7+IL-18, and IL-7+IL-12+IL-18) using monocle3 [10.1038/nmeth.4402]. TCR-seq raw sequencing data were processed using the Cellranger ‘adj’ program. The TCR clonotypes from previously filtered scRNA-seq were used in the further analyses.

### Single-cell RNA-seq data analysis

The preprocessed scRNA-seq data were aligned with the IL-7 treatment group using ‘align_cds’ from the monocle3 program [https://doi.org/10.1038/s41586-019-0969-x], and data were integrated for each set. Each cluster was characterized using the marker gene expression levels and referring to previous studies. The top specifically expressed genes of each cluster were selected by the specificity of each gene, as estimated using ‘FindAllMarkers’ in Seurat v4.0 [https://doi.org/10.1016/j.cell.2021.04.048], and then their expression level was visualized using ‘DoHeatmap’ in the same program. We calculated the percentage of cells in each condition for each cell type by dividing the number of cells in each condition by the number of clustered cells. A DEGs analysis was performed for each cytokine treatment of each set using ‘FindMarkers’ in Seurat with the following parameters: min.pct = 0, min.cells.feature = 1, min.cells.group = 1, logfc.threshold = 0. DEGs between cytokine treatments and DEGs between conditions were defined at FDR <0.05 and average log2 fold change ≥0.1. A functional analysis of DEGs and cluster specific gene sets was performed in MSigDB’s C2 (KEGG) and C5 (GO). The functions were analyzed using the maximum and minimum size of the gene sets (KEGG with min 2 genes and max 500, GO with min 2 genes and max 150 genes). The gene set enrichment test was performed using the ‘enricher’ function of the clusterProfile package [10.1089/omi.2011.0118]. The significance level was set as the function that satisfied FDR < 0.2 and had at least 2 overlapping genes.

### Signature score of the MP CD4^+^ T cells

The signature score of the CD4^+^ T cells was estimated to define the features of each cell type. The monocle3 object of each set of cytokine treatments was transformed into the Seurat object. They were normalized by condition and then integrated. The signature score of each cell type was calculated using AddModuleScore in the Seurat package.

### Trajectory inference of MP CD4^+^ T cells by cytokine stimulation

Effector memory CD4^+^ T-cells have the plasticity to differentiate into various helper cells. We performed a trajectory analysis on the subcluster composed of the initial cells that are likely to be differentiated according to each cytokine stimulus and the cells that responded. First, we selected 2,000 highly variable genes by cell type, and then we inferred the trajectory using reduceDimension with DDRTree in the monocle2 program [10.1038/nbt.2859]. Pseudo-time was calculate using orderCells in the same program. DEGs following the pseudo-time trajectory were defined at a q-value < 1e-50.

### Estimation of regulon activity score

*Regulon* describes the relationship between a transcription factor and its regulatory target genes, and its activity is calculated using the expression levels of the genes in the regulon. First, we constructed a correlation matrix between genes, and then we calculated the activity of each regulon using GENIE3 [10.1371/journal.pone.0012776]. At that time, all the parameters were set to the default. The regulon relationship was derived from mm10__refseq-r80__500bp_up_and_100bp_down_tss.mc9nr and mm10__refseq-r80__10kb_up_and_down_tss.mc9nr in the ‘cisTarget_databases’ of the R package. All processes were performed using the SCENIC program in R [10.1038/nmeth.4463]. Differentially activated regulons were considered only when they contained at least 1% of expressing cells.

### T-cell receptor repertoire analysis

The TCR repertoire of the cells was considered only from the scRNA-seq data. The degree of TCR expansion was categorized as hyperexpanded (100 < X <= 500), large (20 < X <= 100), medium (5 < X <= 20), small (1 < X <= 5), and single (0 < X <= 1). The TCR diversity of each sample was calculated using Shannon’s index, which considers non-uniformity in the frequency of the clonotype (gene and nucleotide sequences). All procedures were performed using scRepertoire [10.12688/f1000research.22139.2] in R.

## SUPPLEMENTARY MATERIALS

## Acknowledgements

We thank Dr. Dongsoo Kyeong and Dr. Younhee Shin (Insilicogen Inc.) for supporting bioinformatic analysis of the scRNA-seq data. We thank Mr. Yeon-Ho Kim and Ms. In Young Song for technical support in the FACS sorting conducted at Hanyang LINC Analytical Equipment Center (Seoul) and the NIH tetramer core facility (Emory University) for providing mouse CD1d PBS-57 (Biotinylated Monomer). We thank Prof. Jeehee Youn (Hanyang University) for kindly providing CD45.1^+^, Foxp3-GFP mice and Prof. Kwang Soon Kim (POSTECH) for helping us purchase germ-free and antigen-free mice.

## Author contributions

M.-Z.C., H.-G.L., and J.-M.C. conceptualized and designed this study. M.-Z.C., H.-G.L. performed and analyzed most of the experiments including bioinformatic analysis. Y.J.L. supported conceptualization for bioinformatic analyses. J.-W.Y., G.-R.K. and J.-H.K. supported experiments. R.T. and B.T.E. provided Bhlhe40^-/-^ and Bhlhe40^GFP^. M.-Z.C., H.-G.L., and J.-M.C. wrote draft manuscript, and all authors reviewed the manuscript. J.-M.C. supervised the analyses and acquired funding.

## Data and materials availability

The data that support the findings of this study are available from the corresponding author upon reasonable request. RNA-seq data have been deposited in the NCBI Gene Expression Omnibus.

## Supporting Information

**Figure S1.**
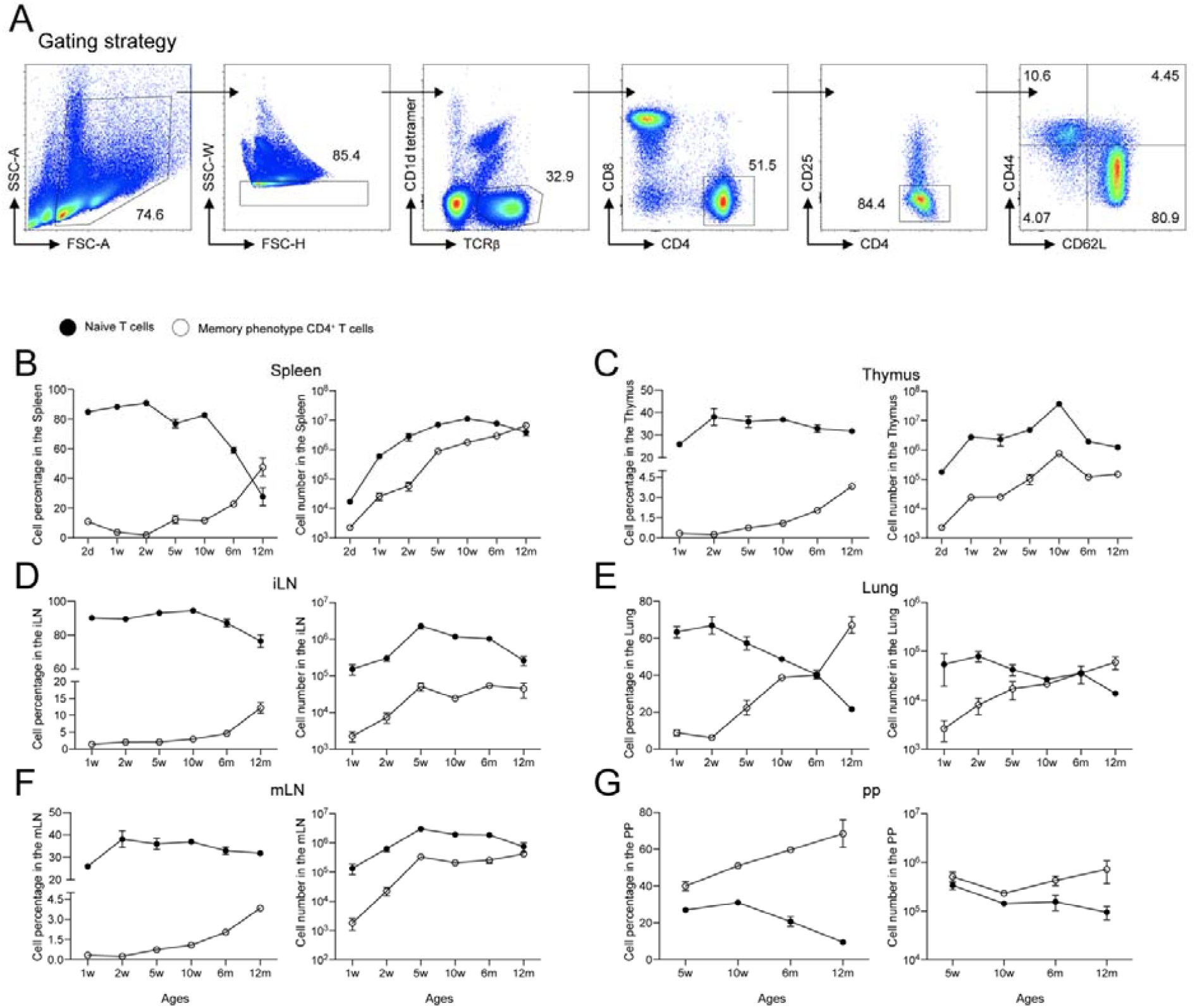
CD44^high^ memory phenotype (MP)^+^ T cells increase with age in mice. (A) Gating strategy of Splenic MP CD4^+^ T cells (TCRβ ^+^ CD1d tetramer^-^ CD8^-^ CD4^+^ CD25^-^CD62L^low^ CD62LCD44)^high^). Analysis of cell percentage and number of naïve (TCRβ^+^ CD1d tetramer ^-^CD8^-^ CD4^+^ CD25^-^ CD62L^low^CD44^high^) and MP CD4T cells (TCRβ CD1d tetramer^-^CD8^-^CD4^+^CD25^-^CD62L^low^CD44^high^) by age from (B) the spleen, (C) thymus, (D) iLN, (E) lung, (F) mLN, and (G) PP.

**Figure S2.**
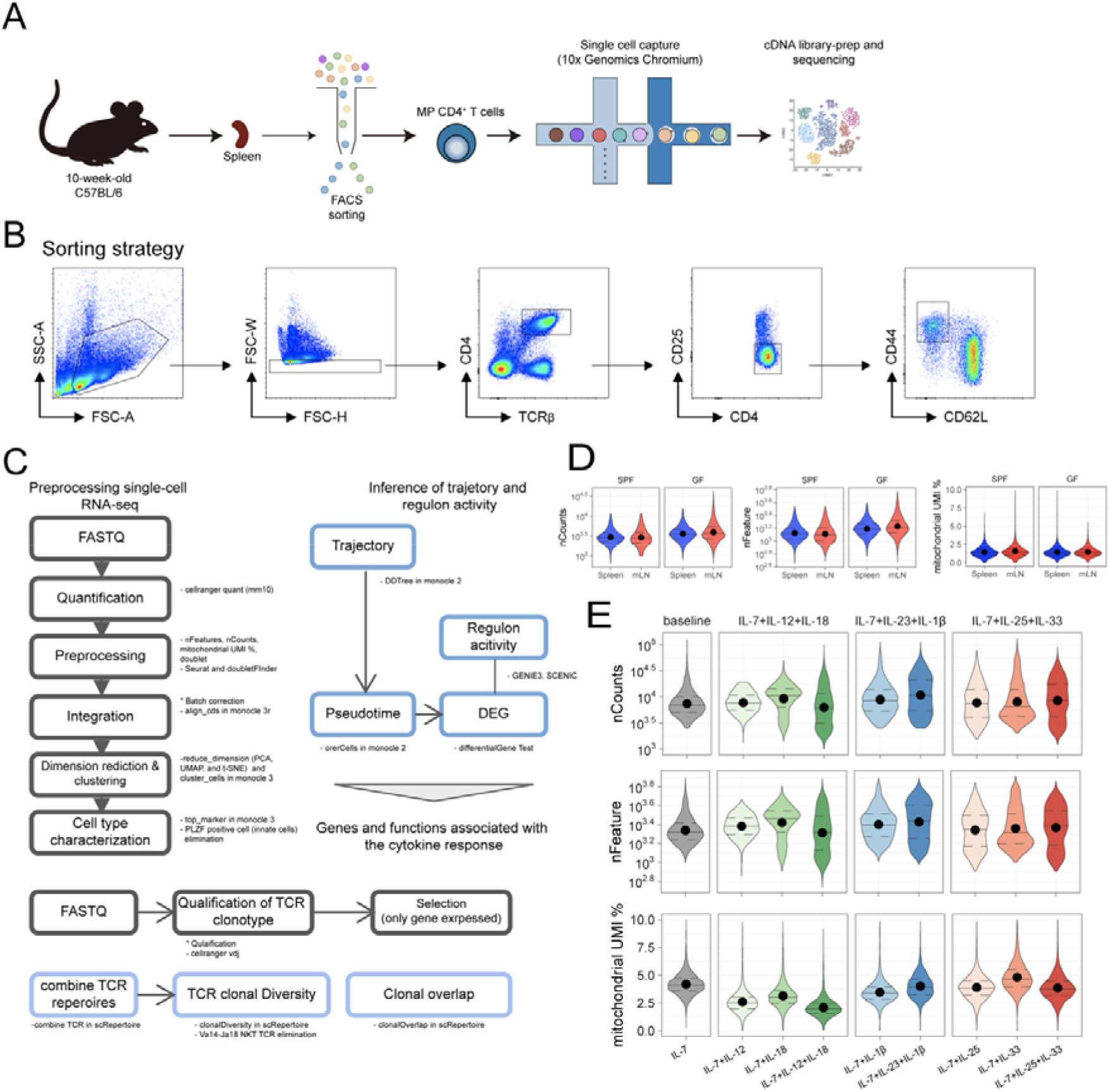
Strategy and flow of single cell RNA-sequencing analysis. (A) Visual concept of scRNA-seq for MP CD4^+^ T cells. (B) Gating strategy of splenic MP (TCRβ^+^CD4^+^CD25^-^CD62L^low^CD44^high^) CD4^+^ T cells. (C) Workflow of scRNA-seq and TCR-seq of PLZF and TCR V_α_14-J_α_18 (TRAV11-TRAJ18) negative MP CD4^+^ T cells. (D) Quality measures of splenic and mLN MP CD4^+^ T cells from SPF- and GF–housed mice. (E) QC metrics for each condition of MP CD4^+^ T cells. (nFeature: the number of expressed genes per cell, nCounts: the number of total UMI counts per cell, mitochondrial UMI %: the percentage of UMI counts of mitochondrial genes per cell)

**Figure S3.**
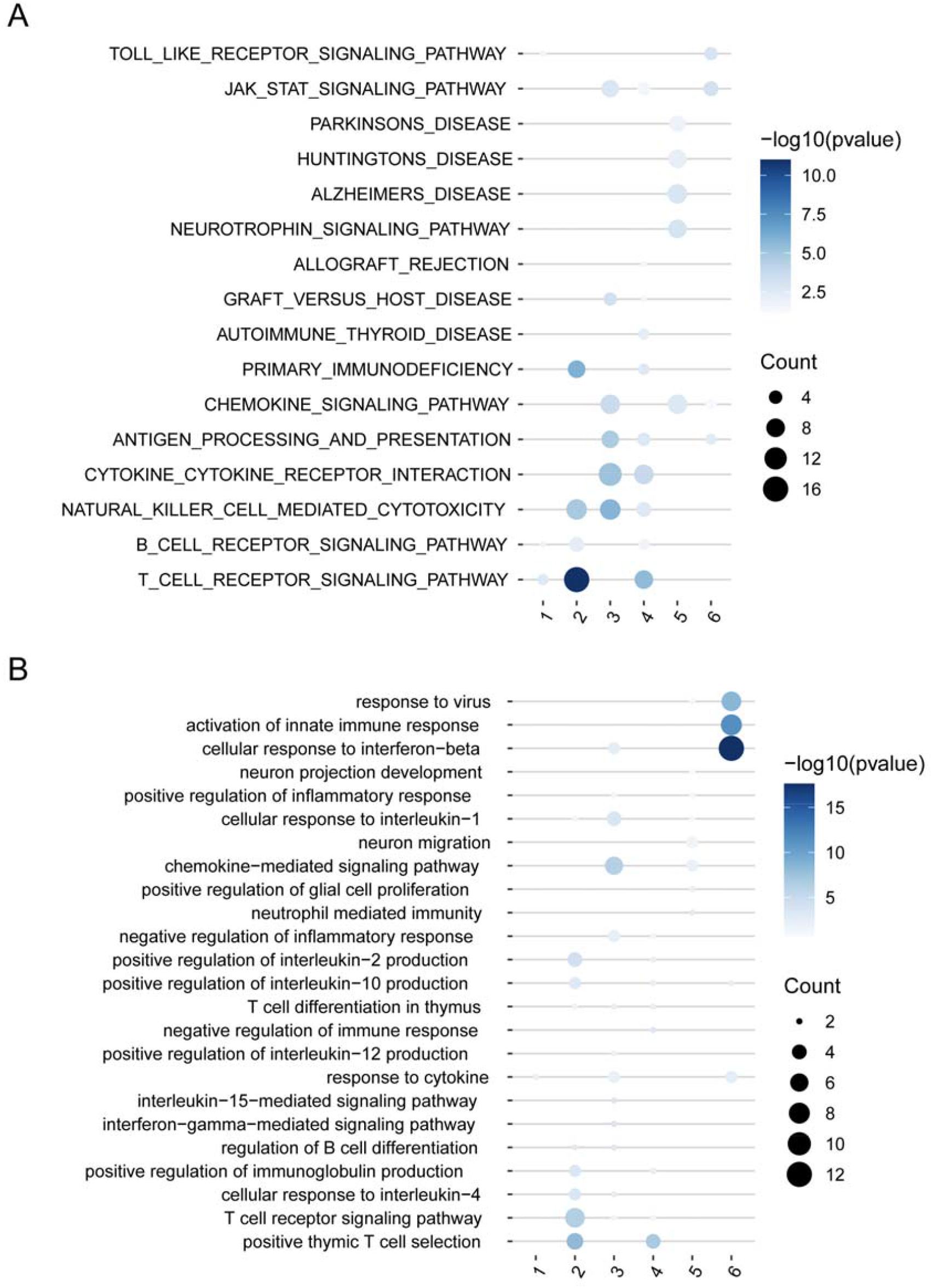
KEGG/GO analysis of steady-state MP CD4 T cells. Selected (A) KEGG and (B) GO terms in each subpopulation of splenic MP CD4^+^ T cells.

**Figure S4.**
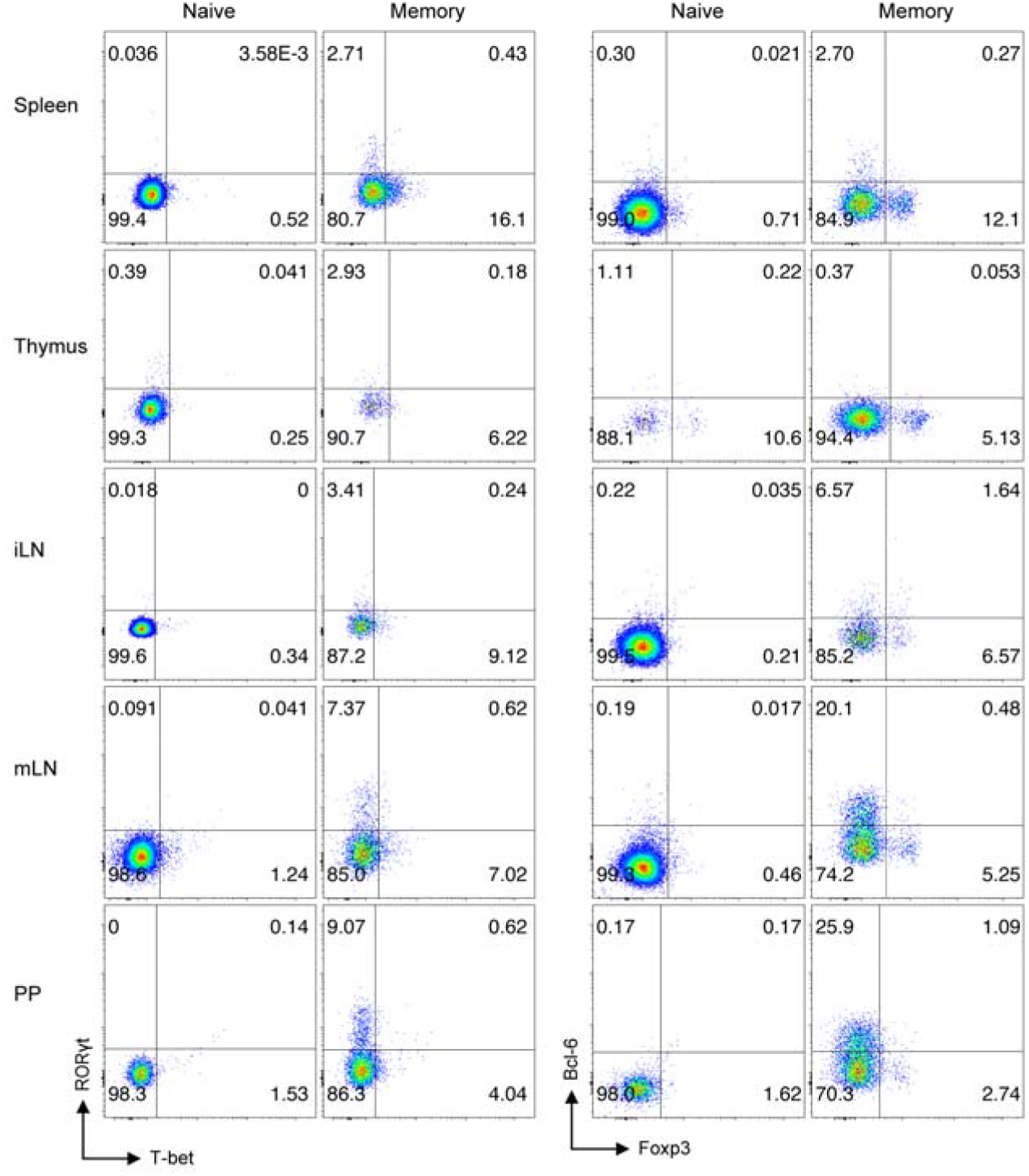
Effector T cell lineage–specific marker expression in MP CD4^+^ T cells. Transcription factor expression levels of T-bet, RORγt, Bcl-6, and Foxp3 in naïve and MP CD4^+^ T cells from the spleen, thymus, iLN, mLN, and PP.

**Figure S5.**
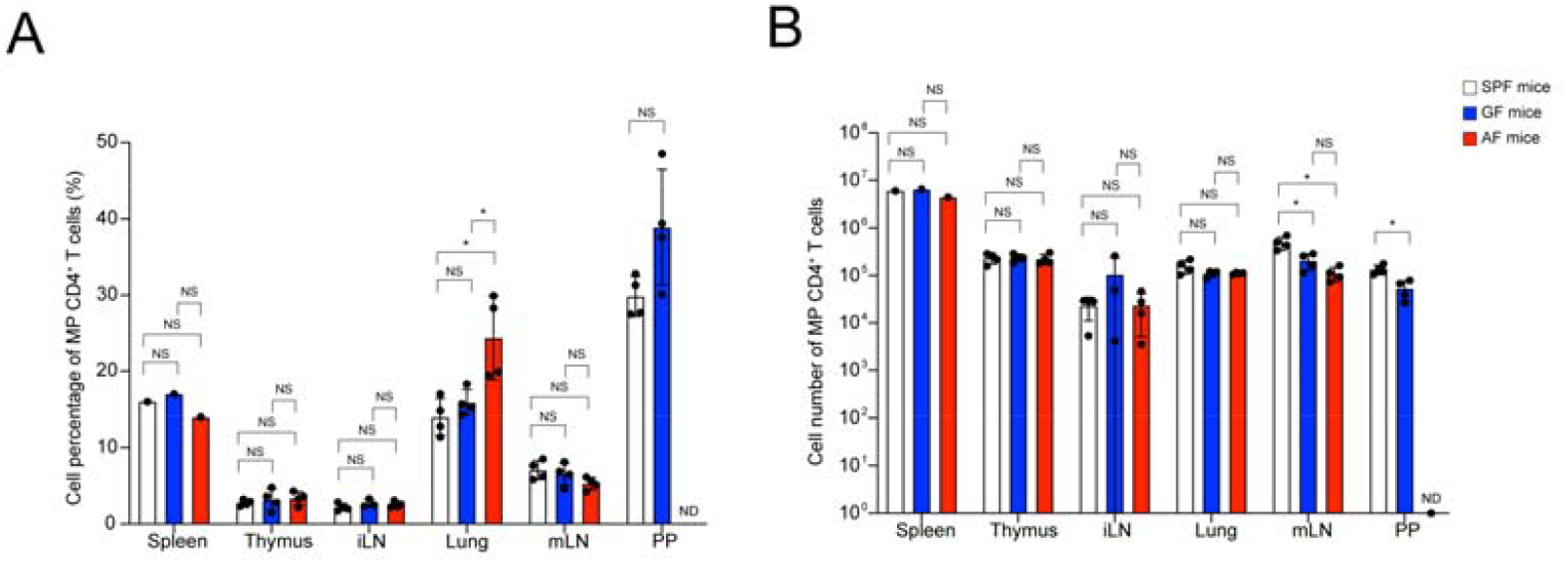
Comparison of MP CD4^+^ T cells in tissues from SPF, GF, and AF mice. (A) Proportion and (B) number of MP CD4^+^ T cells in the spleen, thymus, iLN, lung, mLN, and PP in SPF, GF, and AF mice (n=4). Data are presented as the mean ± S.D. P values were calculated using Mann-Whitney U-test (ND, not detected; NS, not significant; *p < 0.05, **p < 0.01, ***p < 0.001).

**Figure S6.**
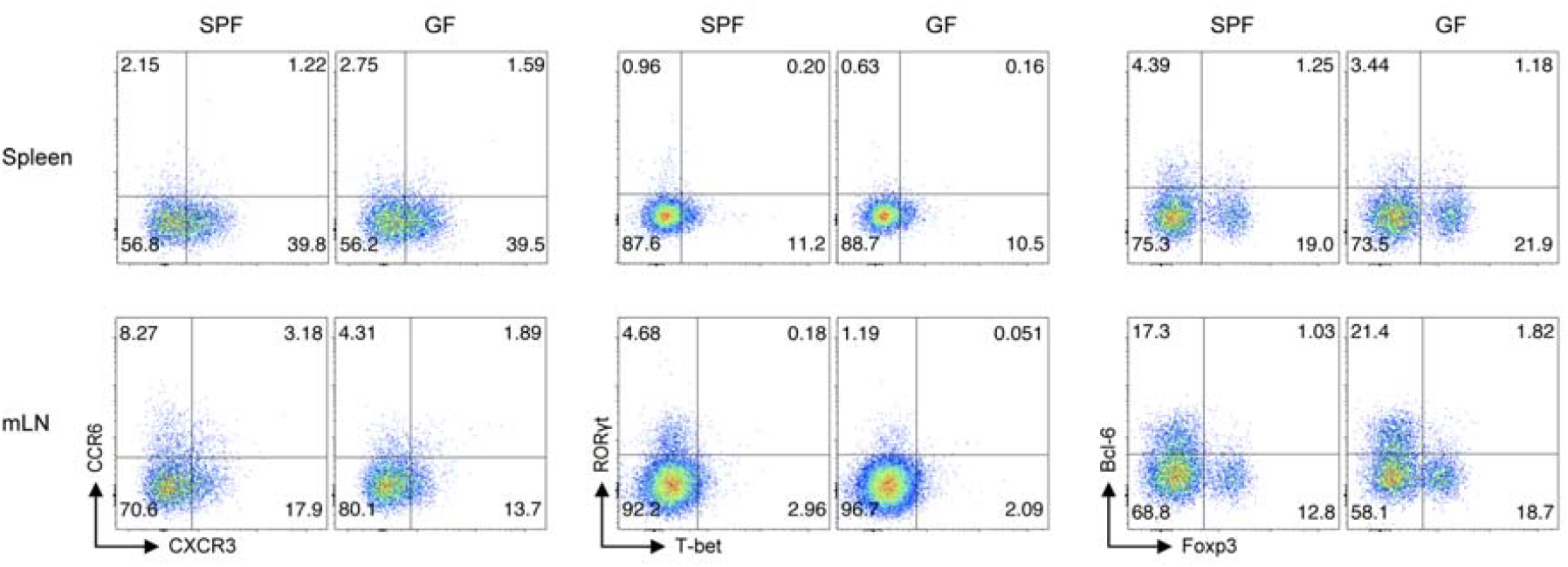
Effector T cell lineage–specific chemokine receptors and transcription factors in MP CD4^+^ T cells from SPF and GF mice. Expression of chemokine receptor of CCR6, CXCR3 and transcription factors of T-bet, RORγt, Bcl-6, and Foxp3 in spleen- and mLN-derived MP CD4^+^ T cells from SPF and GF mice.

**Figure S7.**
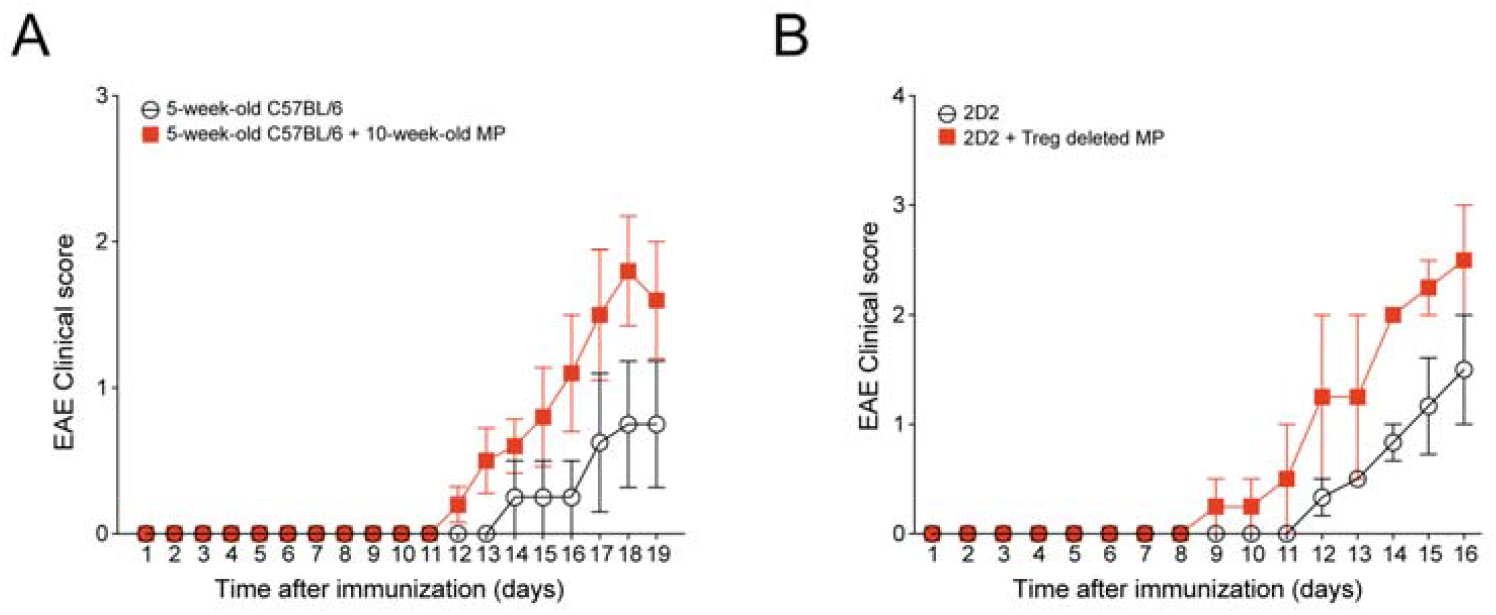
MP CD4^+^ T cells exacerbate autoimmune encephalomyelitis. (A) 5-week-old C57BL/6 mice with or without adoptively transferred 5 × 10^5^ MP CD4^+^ T cells (TCRβ^+^ CD1d tetramer ^-^CD4^+^ Foxp3 ^-^CD62L ^low^ CD44 ^high^) from 10-week-old Foxp3-GFP mice were induced EAE by MOG_35-55_ in CFA (n=4). (B) 5 × 10^4^ naïve CD4^+^ T cells (TCRβ^+^ CD4^+^ V_β_ 11^+^ CD25^-^ CD62L ^low^CD44 ^high^) from 2D2 transgenic mice were adoptively transferred, with or without 5×10^5^ splenic Treg-deleted MP CD4^+^ T cells (TCRβ^+^ CD1d tetramer ^-^CD4^+^ Foxp3^-^ CD62L ^low^ CD44 ^high^) sorted from Foxp3-GFP mice, into Rag^-/-^ mice who were immunized with MOG_35-55_ in CFA (n = 3).

